# An Amber-Encoding Helper Phage for More Efficient Phage Display of Noncanonical Amino Acids

**DOI:** 10.1101/2022.12.19.521047

**Authors:** J. Trae Hampton, Chia-Chuan Dean Cho, Zhi Zachary Geng, Demonta D. Coleman, Peng-Hsun Chase Chen, Gopal K. Dubey, Lauralee D. Sylvain, Shiqing Xu, Wenshe Ray Liu

## Abstract

In the past two decades, phage display has emerged as a powerful technique for the identification of antibodies and peptide ligands for therapeutic targets. Using the amber suppression-based noncanonical amino acid (ncAA) mutagenesis approach, we and others have shown that the chemical space in phage display can be significantly expanded for drug discovery. However, the use of amber codon in phages results in poor phage yields and requires tedious processes to enrich amber codon-containing (amber obligate) phage clones. In this work, we demonstrate the development of a novel helper phage, CMa13ile40, for rapid and continuous enrichment of amber obligate phage clones and efficient production of ncAA-containing phages. CMa13ile40 was constructed by the insertion of a *Candidatus Methanomethylophilus alvus* pyrrolysyl-tRNA synthetase/PylT gene cassette into a helper phage genome. The afforded novel helper phage allowed for a continuous amber codon enrichment strategy for two different phage display libraries and demonstrated a 100-fold increase in selectivity for packaging of library plasmids in comparison with original helper phage plasmids. To demonstrate the applicability of the system, CMa13ile40 was used to create two phage-displayed peptide libraries containing two separate ncAAs, *N*^*ε*^ -*tert*-butoxycarbonyl-lysine (BocK) and *N*^*ε*^ -allyloxycarbonyl-lysine (AllocK), respectively. These were then used to identify peptide ligands that bind to the extracellular domain of ZNRF3, a membrane-bound E3 ligase. Each selection showed differential enrichment of unique sequences dependent upon the ncAA used. Using biolayer interferometry, enriched peptides from both selections were confirmed to have low micromolar affinity for ZNRF3 and this affinity is dependent on the presence of the ncAA used for selection. Our results clearly show that ncAAs in phages provide unique interactions for selection of peptides that are different from each other and from canonical amino acids. As an effective tool for phage display, we believe that CMa13ile40 can be broadly applied to a wide variety of applications.

## INTRODUCTION

Phage display has emerged as a powerful technique for the identification of both antibodies and peptide ligands for therapeutically relevant targets. It typically can be used to screen over 10^9^ ligands for binding to a target at one time, allowing for biopanning of highly diverse libraries. However, it has been limited to express peptides that contain only canonical amino acids (cAAs), severely hindering its application to drug discovery. Drug discovery using peptide libraries with expanded chemistries is an emerging field, with many different campaigns underway for phage display of oligopeptides,^1^ pharmacophore-peptide fusions,^2^ and covalent handles.^3,4^ While these are all exciting advances in the field, they do not allow for high diversity around the areas of expanded chemistry because they utilize cyclic linkers or N-terminal amino acids to functionalize the peptide libraries. Rather, it would give more diverse peptides if expanded functionalities were directly incorporated within the peptide libraries. Genetic code expansion has been successfully done in in vitro experiments that use mRNA display technologies,^5–8^ but has been limited in phage displayed libraries. Recently, our group and others have developed techniques to incorporate noncanonical amino acids (ncAAs) into phage-displayed peptide libraries, which has resulted in two promising techniques: phage display of genetically-encoded cyclic peptides^9,10^ and active site directed ligand evolution.^11^ These strategies utilize a three-plasmid system to incorporate ncAAs into phage libraries that includes a helper phage plasmid, a plasmid to encode the library, and a plasmid for suppression of amber codons with ncAAs (Figure 1). Alongside this, our group has developed a method that takes advantage of superinfection immunity to enrich phages that contain amber codons within library regions, which was essential to prevent enrichment of parasitic sequences that lack amber codons but are more productive in propagation.^11^ While these systems are revolutionary in their ability to encode ncAAs into phage libraries, they result in much lower phage yields in comparison with phages that only contain 20 cAAs. This further limits the diversity of phage libraries that contain ncAAs, thus hindering the selection potential of these libraries. Therefore, new strategies to express and enrich amber codon-containing (amber obligate) clones are needed to further develop phage libraries that contain ncAAs.

**Figure 1:**
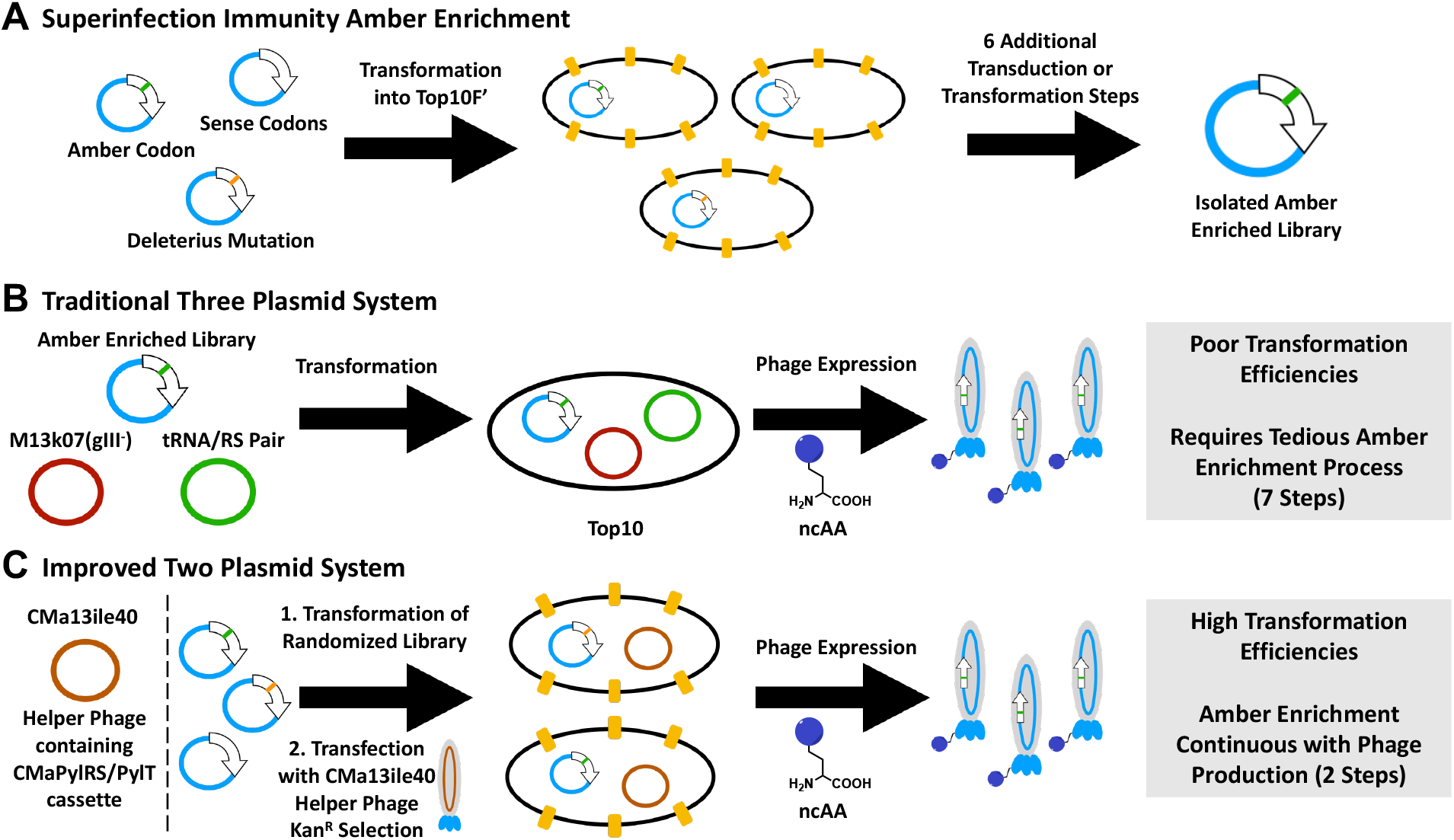
Concept for novel helper phage design. A) Previously developed amber enrichment process involves transformation of the randomized library into Top10F’ followed by 6 iterative transduction and transformation steps. B) The traditional three-plasmid expression system for expression of phage libraries containing ncAAs. A previously amber enriched library is transformed into Top10 cells along with two plasmids for expression of phages. C) The two-plasmid system that contains the novel helper phage that allows for both amber enrichment and phage expression simultaneously.

In the past two decades, different efforts from our group and others have allowed for genetic code expansion through amber suppression using more than 200 different amino acids.^12–15^ The first studies for amber suppression in phage display took advantage of orthogonal aminoacyl-tRNA Synthetases that were derived from either *Methanocaldococcus jannaschii* (MjTyrRS)^10^ or from *Methanosarcina mazei* (MmPylRS) and their corresponding amber suppressing tRNAs.^9,11^ Recently, a novel PylRS/PylT pair, CMaPylRS/PylT, was discovered in *Candidatus Methanomethylophilus alvus*, a methanogenic archaea found in the human gut.^16,17^ In comparison to MmPylRS, CMaPylRS is much smaller (31 kDa) and contains a shortened *N*-terminal domain that leads to increased solubility and stability of the protein.^18,19^ Like with the originally discovered PylRS strains, multiple groups have begun campaigns to repurpose the CMaPylRS for encoding of ncAAs or to improve its selectivity/orthogonality in comparison to other PylRSs.^18,20–24^

We looked to take advantage of the truncated structure of CMaPylRS to improve the efficiency of genetic code expansion in phage display. Because of its small size, we envisioned that the CMaPylRS/PylT pair could be easily integrated into the helper phage plasmid and allow for efficient packaging of the helper phage. In this study, we report the development of a novel helper phage that contains a CMaPylRS/PylT gene fragment and take advantage of this to development an optimized amber enrichment strategy for phage display of ncAAs.

## RESULTS

### Rationale and Design of Amber-Supressing Helper Phage

We have previously described a system for the creation of amber obligate peptide libraries.^11^ While this technique is successful in preventing the overexpression of parasitic sequences that contained only cAAs, it requires several tedious transformation and transduction steps, each of which could possibly lead to bias and reduce sequence diversity (Figure 1A). Alongside this, after the amber enrichment process, a three-plasmid system is necessary to produce phage displayed peptide libraries containing ncAAs (Figure 1B). Because this system requires transformation of the amber-enriched plasmid into cells containing two other plasmids for phage production (one to produce phage proteins and another for amber suppression), it is also hindered by poor transformation efficiencies that limit the initial library coverage to 10^9^ total peptides. We envisioned a way to enhance the amber enrichment and phage production processes would be through the development of an amber suppressing helper phage (Figure 1C) that fuses the two accessory plasmids from the initial three-plasmid system. This would allow for a continuous amber enrichment and phage expression process that eliminates five transformation and transduction steps. Also, it would allow for efficient transformations of the initial library plasmid into cells that do not contain any other plasmids, resulting in the ability to cover larger library sizes. Because of these potential advantages, we looked to create a helper phage that was able to perform amber suppression.

### Development of Helper Phage CMa13ile40

To develop a novel helper phage for incorporation of ncAAs into phage displayed peptides, we first looked for a good site for insertion of the CMaPylRS/PylT gene cassette into the helper phage DNA. Currently, two different styles of helper phages are used for phage display. One of these contains a full-length pIII protein that allows for simultaneous display of both wild-type pIII and pIII modified with the peptide library.^25^ Although this may be helpful in avoiding any valency effects during the selection process, typically the presence of wild-type pIII within the helper phage results in little to no displayed peptides containing ncAAs. This is due to the gIII gene from the helper phage that contains only sense codons outcompeting the production of pIII that is dependent upon amber suppression for translation. Because of this, we looked to use a helper phage that was deficient of pIII production. Recently, we reported using a helper phage plasmid (M13k07TAA) that contains a stop codon (UAA) within gIII to prevent expression of wild type pIII.^11^ However, for our purposes, we envisioned CM13d3 (Antibody Design Labs), an M13 phage that consists of almost a fully deleted gIII and an interference-resistant mutation, would be a better vector for packaging due to its truncated size.^26,27^ Even with the inserted CMaPylRS/PylT gene cassette in CM13d3, the helper phage genome would be similar in size to wild-type helper phage, so we believed this would be more likely to efficiently package into virions. Therefore, we first designed CMa13d3, a construct that contained the CMaPylRS/PylT gene cassette within the intergenic DNA of CM13d3 (Figure 2A). To test the packaging efficiency of the helper phage, we cotransformed it with a plasmid expressing gIII (pBAD-gIII), which does not contain the proper f1 origin for packaging, and expressed pIII to allow for packaging of the helper phage. However, the modification resulted in poor phage yields (10^8^ pfu/mL, Figure 2B) that were not desirable for use as a helper phage. We believed this was due to a disruption in the packaging efficiency of the plasmid. Helper phage plasmids contain 100-fold decreased packaging efficiency due to a split f1 origin, which is required for recognition by pII to create single-stranded DNA to be packaged into virions.^28–30^ In CMa13d3, we have increased the distance between the two sites of the f1 origin, which may disrupt the recognition by pII even more. Also, recognition by pII is dependent upon the superhelicity of the DNA, which may have been hindered by the large insertion into the intergenic DNA.^31^ Therefore, we next tried to modify the plasmid to increase phage production of our new construct. To potentially increase the production of pII, we deleted the operator (CMa13opdel) that controls transcription of gII. Also, we created another construct (CMa13ile40) the contained a Met40Ile mutation in pII, which has been previously reported to increase the cooperativity of the enzyme and allow for enhanced recognition of DNA.^29,32^ Both mutations gave significantly increased phage yields, with CMa13opdel increasing production 3-fold and CMa13ile40 increasing it 8-fold (Figure 2B). We deemed the phage yields of CMa13ile40 to be high enough for performing future experiments involving phage display.

**Figure 2:**
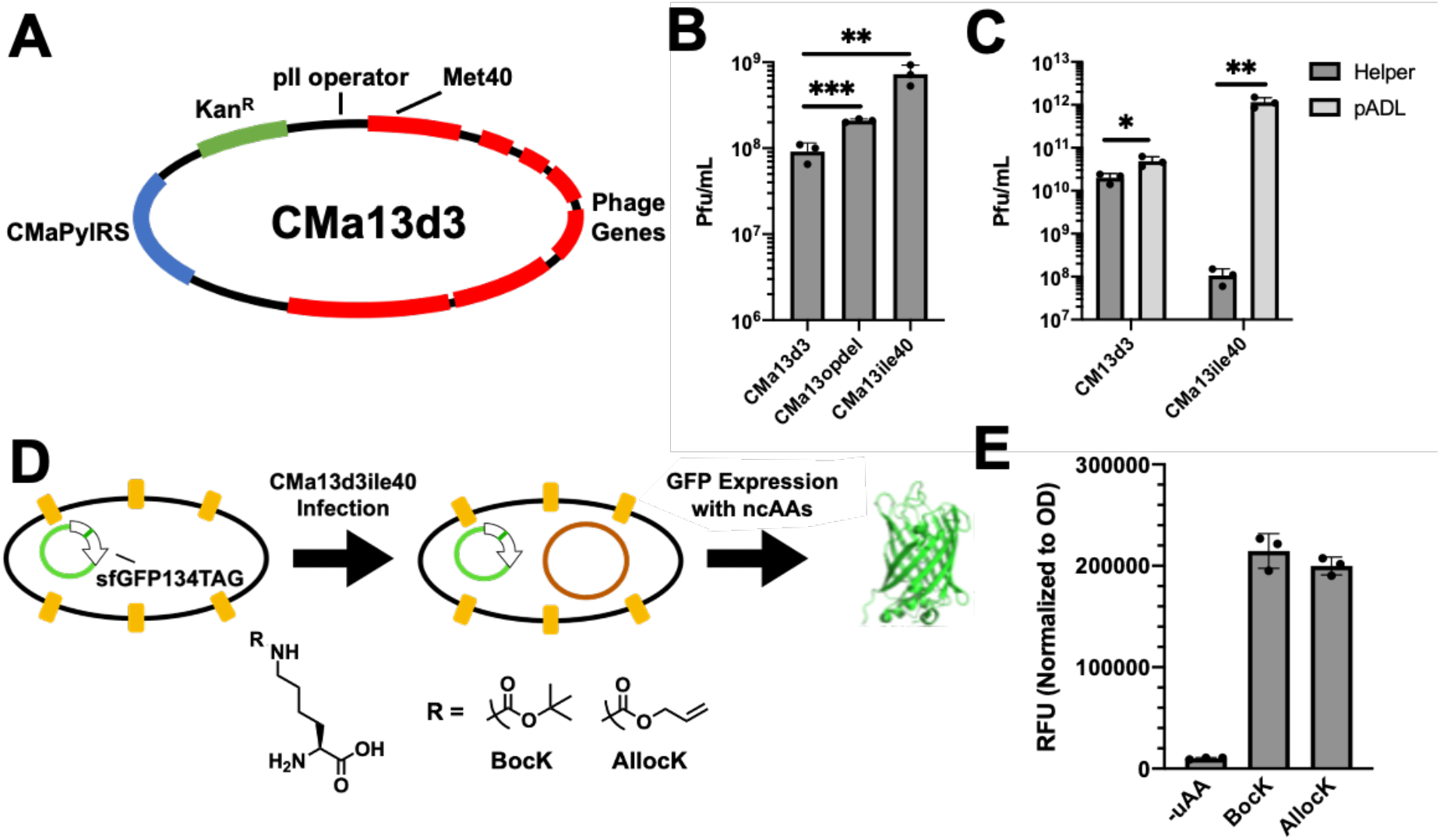
Design of helper phage plasmid and subsequent validation of CMa13ile40. A) Initial plasmid map for CMa13d3 with sites highlighted where mutations were inserted (pII operator and Met40). B) Helper phage expressions of the different mutants generated. Opdel: pII operator deletion, ile40: Met40Ile mutation in pII. Phage titers are plotted as mean ± s.d. for n = 3 biological replicates. C) Packaging selectivity assays for expression of the helper phage vs. a pADL-AAKAA plasmid. Phage titers are plotted as mean ± s.d. for n = 3 biological replicates. D) Experimental design for testing incorporation of ncAAs into sfGFP using CMa13ile40. E) GFP fluorescence measurements using 1 mM BocK, 1 mM AllocK, or no ncAA (-uAA). All experiments are displayed as mean ± s.d. for n = 3 biological replicates.

### Amber Suppression using CMa13ile40

Before using CMa13ile40 to perform phage display of ncAAs, we looked to validate the new helper phage to ensure everything functioned as desired. Because the modification in pII affects the promiscuity of the enzyme for supercoiled DNA, we wanted to ensure that this did not result in poor selectivity when packaging the phage libraries. To test this, we performed a mock phage expression using a plasmid that contained unmodified gIII and a full f1 origin for packaging (pADL-AAKAA) and expressed these phages using either CM13d3 or CMa13ile40 as helper phage. To quantify the selectivity of phage production, the phages were titered separately on ampicillin and kanamycin containing plates, since the helper phage have kanamycin resistance and the pADL plasmid has ampicillin resistance. Phages produced using CMa13ile40 demonstrated 10,000-fold higher selectivity for packaging the pADL plasmid than the helper phage plasmid (Figure 2C). On the other hand, phages produced using CM13d3 only displayed about 2.5-fold higher selectivity in packaging the pADL plasmid (Figure 2C). Also, pADL-containing phages produced by CMa13ile40 were approximately 20-fold higher concentration than those produced by CM13d3. From this, we can infer that CMa13ile40 is a proficient helper phage for phage display and exceeds the current standards set by CM13d3. To ensure that CMa13ile40 could efficiently incorporate ncAAs at stop codons following its infection into cells, we transformed a plasmid that expresses GFP with a D134TAG mutation (pBAD-sfGFP-134TAG) into Top10F’ cells, which contain the F pilus to allow for phage infection. After infecting the cells with CMa13ile40, GFP production was then induced with or without the presence of ncAAs (Figure 2D). GFP fluorescence was then measured in the cell lysates after expressing for 24 hours, and the fluorescence was normalized to the OD_600_ of the cultures to account for any differences in cell growth between replicates. Similar to previously reported results using CMaPylRS, we observed high GFP production only when supplementing cells with 1 mM of either *N*^*ε*^ -Boc-Lysine (BocK, Figure 2E) or *N*^*ε*^ -Alloc-Lysine (AllocK, Figure 2E). Without the addition of either ncAA, the fluorescence was 200-fold lower, indicating a lack of amber suppression from cAAs alone. Considering both these results and the phage packaging data, we believed CMa13ile40 would be effective as a helper phage for the incorporation of ncAAs into peptide libraries.

### Amber Enrichment of Phage-Displayed Peptide Libraries Using CMa13ile40

Following validation of the CMa13ile40 helper phage for phage display and amber suppression, we then looked to test its ability to perform amber enrichment of a phage library. Previously, we have taken advantage of superinfection immunity to enrich amber obligate phages.^11^ However, this process is very tedious, lasting multiple days and requiring multiple transformation/overnight growth steps that could lead to amplification of parasitic sequences (Figure 1A). Using CMa13ile40, we theorized that five steps of the process could be removed since the amber enrichment and phage expression with an ncAA could be performed in the same day (Figure 3A). To test this, we transformed a phage library containing Cys(NNK)_12_Cys, where NNK is a reduced codon set including amber codon, on the *N*-terminal domain of pIII into Top10F’. To enrich for clones that contained amber codons, we then induced expression of pIII without adding any ncAA. In theory, this would lead to production of pIII only in clones that do not have amber codons, which prevents these clones from being infected by another phage. Following induction of pIII expression, the cells were infected with CMa13ile40, then resuspended in media containing kanamycin, 1 mM IPTG, and 5 mM BocK to produce phages. To ensure that the amber enrichment was successful, we employed several different controls: 1) A control where pIII was induced before infection but without BocK in the final media; 2) A control where pIII was not induced before infection, but BocK was included in the media; 3) A control where pIII was not induced before infection, but BocK was excluded in the final media. After an overnight expression, the phages were purified and titered to estimate the amber enrichment (Figure 3B). In the experiments that included the induction of pIII prior to infection, we observed a 100-fold difference between the phage yields that was dependent on the presence of BocK. However, without the induction, there was no dependence on BocK for phage expression.

**Figure 3:**
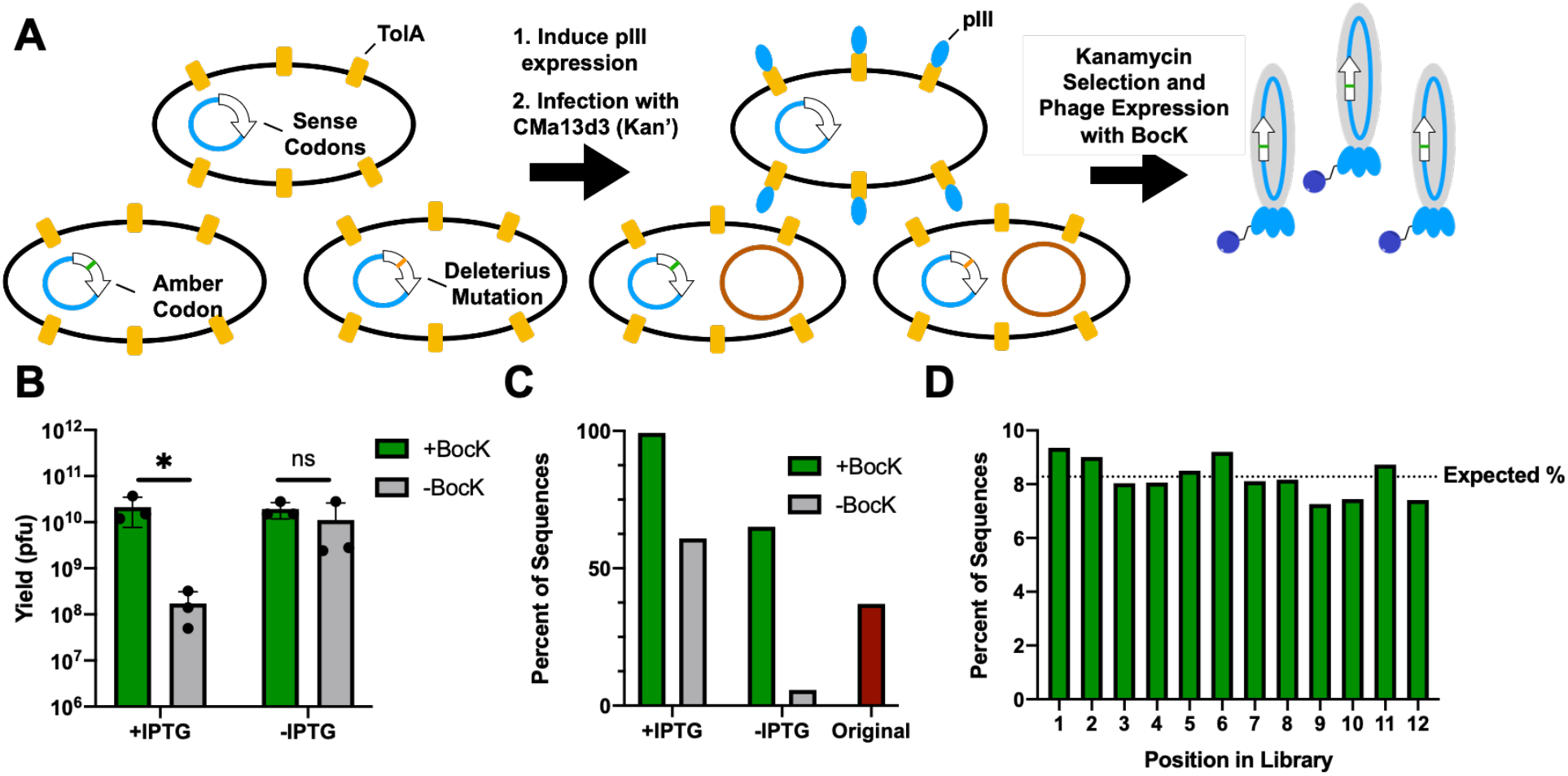
Amber enrichment of a 12-mer phage library using the novel helper phage CMa13ile40. A) Experimental design of the amber enrichment and phage expression process using BocK. B) Phage titers after expression with or without BocK for the amber enriched clones (+IPTG) vs. clones that lacked the initial pIII expression step (-IPTG). Phage titers are plotted as mean ± s.d. for n = 3 biological replicates. C) Illumina next generation sequencing results of the phages that were produced in section B. The percent of sequences refers to the percent of sequences that contain at least one TAG within the library region. The original TAG content is shown in red. D) The distribution of the amber codon in the amber enriched library. The percent expected if the library was truly randomized for TAG content is shown as a dashed line.

Initial phage titers indicated that the amber enrichment process was successful and that the CMa13ile40 was effective in producing phages that contained BocK. We next looked to sequence the enriched libraries to ensure that the amber enrichment occurred successfully without inducing bias. To do this, we isolated the phage DNA and prepared it for Illumina next generation sequencing through two subsequent PCR amplification steps to amplify out the library region and append adaptor tags to each. The amplicons were then sequenced (MiSeq, 2x150 bp, paired-end). In a similar manner to previously reported works, we used paired-end processing to filter the reads and ensure no sequencing errors or poor quality affected the data. As expected, strong enrichment of amber codons was observed only in the experiment that induced pIII expression prior to infection and contained BocK in the final media. Out of 126,983 reads, 99.8% of them contained at least one amber codon in the library region (Figure 3C). To ensure that the original library was not biased in any way, we also sequenced it and observed that only 37% of the clones contained an amber codon (Figure 3C), which is consistent with the expected percentage (32%) based on NNK randomization. It could be possible that the incorporation of BocK is favored toward to particular sequences, leading to bias within the amber-enriched library. Heat maps of the amino acid distribution at each position indicated that the amber enrichment did not lead to strong bias (Figure S1) and the enriched library showed a distribution of the amber codon throughout the twelve possible positions that would be similar to a randomized distribution (Figure 3D). The 12mer library had a relatively large percentage of clones (37%) that contained amber codons initially. To see if our method would also work with smaller libraries, we tested the amber enrichment using a Cys(NNK)_5_Cys (5mer) library, which initially only contains 15% of clones with at least one amber codon. We also observed high amber enrichment with this library, with 93.4% of sequences (58,273 total) containing at least one amber codon after one cycle of amber enrichment using BocK with CMa13ile40 (Figure S2). This amber content was similar to a previously enriched 7mer peptide library (94% amber-containing clones) that used the old strategy for amber enrichment and has been used for multiple successful phage selections in our lab.^11^ Because there was enrichment of the rare arginine codon (AGG) in the non-amber containing sequences, we hypothesize that the remaining clones consist of low-expressing sequences that are composed of rare codons (Figure S3). Altogether, these data indicate that CMa13ile40 and the improved amber enrichment selection strategy are validated tools for accelerated amber enrichment of phage-displayed peptide libraries.

### Selection of Peptide Ligands for the Extracellular Domain of ZNRF3

After validating the CMa13ile40 helper phage for amber enrichment, we then looked to test its ability to perform a selection using ncAAs. Because the helper phage incorporated both AllocK and BocK efficiently into GFP, we hypothesized that we would be able to use the same helper phage and library to identify different peptide ligands by simply varying the ncAA used during the selection. To do this, we expressed phages with the 12-mer enriched library using both BocK and AllocK. During each phage expression, we monitored the difference between phage yields with and without ncAA to ensure that the expression was dependent upon ncAA incorporation. Each expression showed at least a 10-fold higher phage yield when ncAA was present compared to a negative control without ncAA (Figure S4). We then used the expressed phages to select for peptides that bound to the extracellular domain of ZNRF3, a membrane-bound E3 ubiquitin ligase that has been used to create proteolysis-targeted chimeras (PROTACs) for degradation of oncoproteins.^33^ Each selection was performed in duplicate to ensure reproducibility. After 3 rounds of selection, we observed strong enrichment of phages that bound to the protein target, which was indicated by the percent of phages eluted at the end of the selection (Figure 4B). These enrichment profiles were consistent across the two replicate selections and varied depending upon the ncAA used in selection. We then performed Illumina sequencing of the isolated libraries, using paired-end sequencing analysis as previously described. For each selection, there was strong enrichment of consensus sequences (Figure 4A, 4C, 4F). To ensure the selected peptides were significantly enriched, we made volcano plots for each of the selections by combining the two replicates (Figures 4C and 4F). For the AllocK selections, one peptide accounted for an average of 89% of all the sequences after the third round of selection, and showed a p-value less than 0.05, indicating significant enrichment (Figures 4A, 4C, and S5). In comparison, the BocK selection showed slightly more complexity, but two peptides were identified based on the volcano plot and total sequence percentages in the third round of selection (Figures 4F, 5C, and S6). The second most abundant peptide was determined to not be statistically significant between the two selections, as there was only strong enrichment in the second replicate, resulting in a p-value greater than 0.05. Therefore, using the selection results, two peptides (ZPB1 and ZPB2) were identified to be significantly enriched during the BocK selection. Also, both selections gave completely different heat map profiles of the amino acid abundance at each position, further confirming the distinctive effects of using different ncAAs on selection results (Figure 4D, 4G).

**Figure 4:**
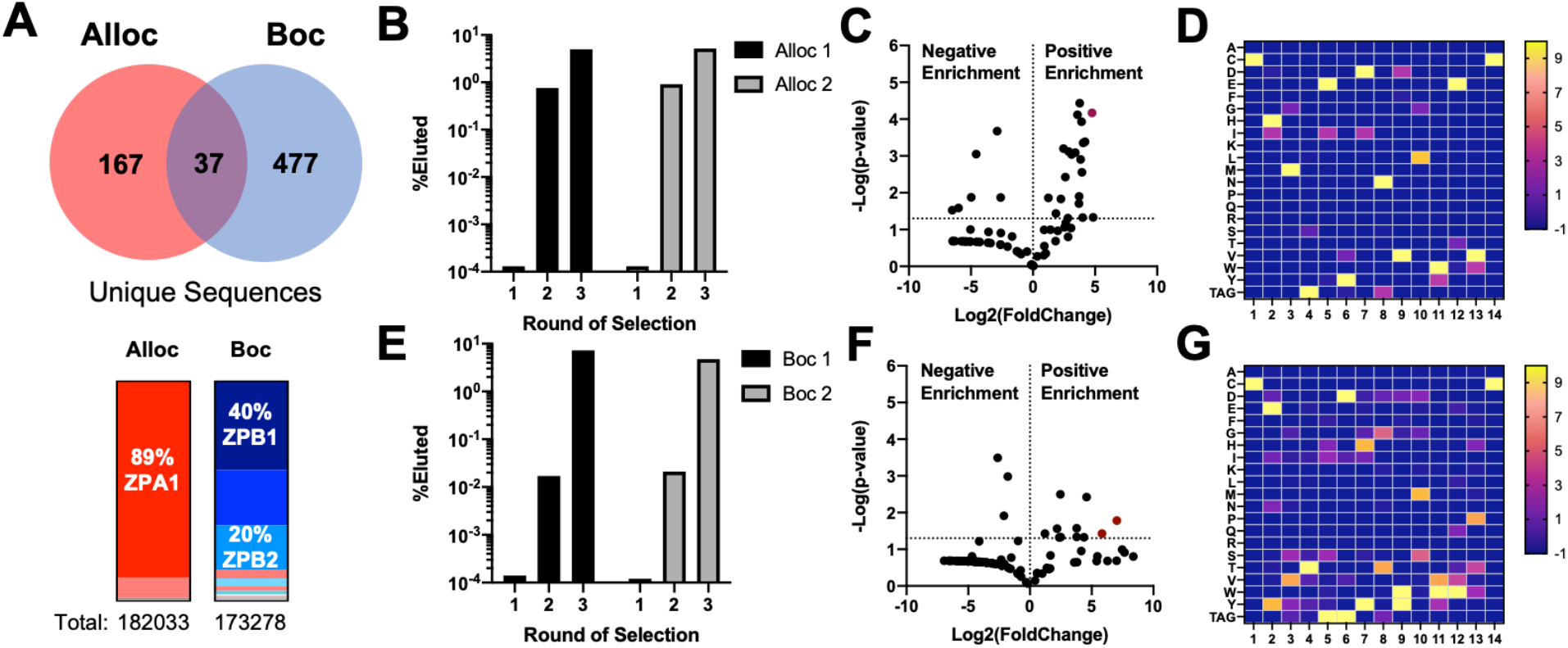
Summary of selection results against ZNRF3 using libraries containing BocK and AllocK. A) Overview of selection results. The Venn diagram corresponds to the total unique sequences found by Illumina sequencing of the third round of each selection (n = 2 for each ncAA). The bars for each correspond to the average percentages of each peptide identified in the selections (n = 2 for each ncAA). Regions in red correspond to those found in the AllocK selections alone, while blue correspond to the BocK selection alone. Pink regions are sequences that were found in both BocK and AllocK selections. B) Phage elution percentages for each round of selection with both AllocK replicates. (%Eluted = Eluted/Input x 100) C) Volcano plot for the peptides identified in the AllocK selections against ZNRF3. The dashed line for p-value corresponds to p = 0.05. The highlighted data point corresponds to ZPA1. D) Heat map for the peptides identified after three rounds of selection using AllocK. E) Phage elution percentages for each round of selection with both BocK replicates. F) Volcano plot for the peptides identified in the BocK selections against ZNRF3. The dashed line for p-value corresponds to p = 0.05. The highlighted data points correspond to ZPB1 and ZPB2. G) Heat map for the peptide library after three rounds of selection using BocK.

While we believed the selection results helped to validate the CMa13ile40 helper phage for use in phage selections involving different ncAAs, we looked to further confirm binding of the peptides to ZNRF3 through biophysical assays. Using solid-phase peptide synthesis, we synthesized the enriched peptides and oxidized them to create cyclic peptides through disulfide formation. These should be the dominant form during selection, as displayed peptides get oxidized in the periplasmic space during packaging. After purification, all peptides were characterized using high resolution mass spectrometry (Figures S10-S16, Table S1). Using biolayer interferometry (BLI), we tested the binding of the peptides to ZNRF3. The top peptide from the AllocK selection, ZPA1, showed low micromolar affinity with a K_D_ of 5.9 ± 0.9 µM (Figure 5B). As a control, we also synthesized the peptide substituting lysine at the AllocK position (ZPA1K), and it exhibited 17-fold lower affinity for ZNRF3 with a K_D_ of 100 ± 10 µM. To test if the linear peptide was binding, we replaced the cysteines with an aminobutyric acid analog at the cysteine positions (ZPA1Abu), and this peptide exhibited no significant binding up until 40 µM of peptide (Figure 5A, S7). For the BocK selection, two of the top peptides, ZPB1 and ZPB2, were synthesized and characterized for their binding using BLI. Both gave similarly low micromolar affinities for ZNRF3, with ZPB1 and ZPB2 having K_D_ values of 4.5 ± 0.9 µM and 1.5 ± 0.4 µM, respectively. To elucidate whether the BocK was essential for binding, we also synthesized free lysine derivatives, ZPB1K and ZPB2K. ZPB1K did not show any binding to ZNRF3, while ZPB2K showed slightly weaker affinity with a K_D_ of 4.8 ± 0.2 µM. This indicates that for ZPB1, the BocK residue is essential for binding. On the other hand, it plays less of a role for binding with ZPB2, possibly with the BocK residue being solvent exposed. Future crystallographic studies and alanine scanning will be used to further elucidate the roles of each residue in binding and potentially allow for further optimization of the peptides. Nevertheless, from these results we can conclude that the CMa13ile40 helper phage is effective in incorporating ncAAs into a phage-displayed peptide library and in identifying unique ligands for target proteins that are dependent upon used ncAAs.

**Figure 5:**
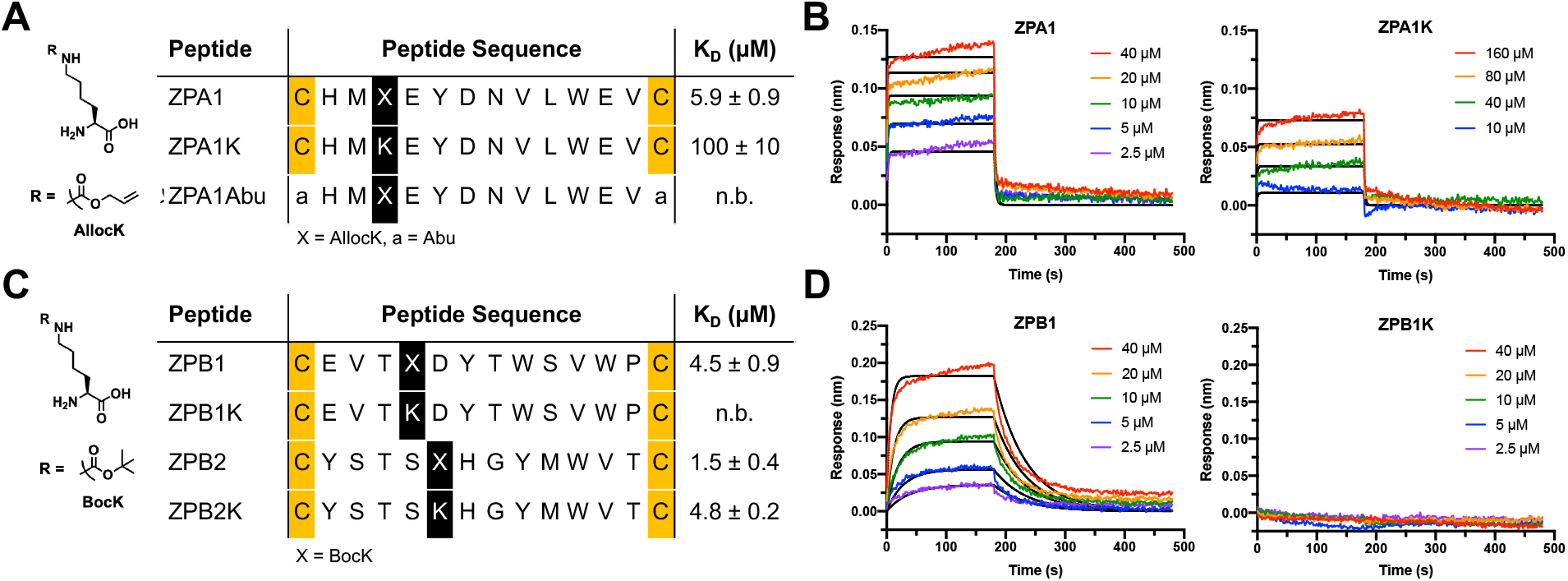
Validation of peptides identified in the selections using BocK and AllocK. A) Peptide sequences from the AllocK selection that were tested for binding to ZNRF3 using biolayer interferometry. K_D_ values are reported as mean ± s.d. of three biological replicates. B) Representative biolayer interferometry traces for peptides from panel A. Data was fit using a 1:1 binding model (black). C) Peptide sequences from the BocK selection that were tested for binding to ZNRF3 using biolayer interferometry. D) Representative biolayer interferometry traces for peptides from panel C. Data was fit using a 1:1 binding model (black).

## DISCUSSION

In summary, we have developed a novel helper phage, CMa13ile40, which contains the CMaPylRS/PylT gene cassette to simplify genetic code expansion in phage display, and we have taken advantage of this helper phage to identify ligands for ZNRF3. Given the E3 ligase activity of ZNRF3, we envision these ligands can be used in future PROTAC experiments to downregulate therapeutically relevant membrane proteins. There are many advantages that this phage expression system has over the traditional three-plasmid approach for phage display of ncAAs. The first of these relates to the efficiency of the initial library transformation. Previously, it was necessary to cover a large library size by transforming into cells that contain two plasmids. This leads to difficult transformation conditions and low efficiency, hindering the size of libraries that can be covered. Also, to perform the original amber enrichment process (Figure 1A), three different transformation steps are required. This significantly hinders the speed to produce phage libraries and could result in significant bias for sequences that transform better than others. However, with this new method, only one initial transformation is necessary into Top10F’, allowing for facile coverage of large libraries (>10^10^ transformants). We also demonstrate increased packaging selectivity of pADL-10b libraries using CMa13ile40, which displayed at least 100-fold better selectivity in comparison to the CM13d3 helper phage. This allows for less interference by the helper phage DNA during the phage display process. We believe this is due to a reduction in the packaging efficiency of the helper phage as a result of the insertion in the intergenic DNA. In addition to validation of the CMa13ile40, we have demonstrated its versatility by performing a selection against the extracellular domain of ZNRF3 using two different ncAAs, BocK and AllocK. The use of two different ncAAs allowed for the identification of two completely different peptides from the same original peptide library, which indicates that specific ncAAs play major roles in the selection process. While we have demonstrated the ability to incorporate both BocK and AllocK into phage libraries, it would be beneficial to have the ability to encode for more diverse ncAAs. We have previously used ncAAs to direct phage selections for active sites of epigenetic readers. However, these studies require the ability to incorporate lysine derivatives other than BocK and AllocK. Although CMaPylRS is still limited in its ability to incorporate diverse ncAAs, the amber enriched phage libraries can also be used to infect Top10F’ and used with the original three plasmid approach with more diverse PylRS/PylT pairs. There have been several different campaigns to begin diversifying CMaPylRS so that it can accept other ncAAs. Recently, a study using phage-assisted non-continuous evolution (PANCE) to evolve the enzyme to more efficiently incorporate BocK was reported.^21^ Because we are able to produce CMa13ile40 phages, we envision the ability to evolve the helper phage to incorporate other ncAAs using PANCE. We are currently exploring this potential, as it would greatly enhance the discovery of new CMaPylRS variants and could be used in a variety of genetic code expansion applications outside of phage display.

## METHODS

### 1. Primer Information and Plasmid Construction

#### Primer List

1. pII-ile40-for: 5’-CGGCATTAATTTATCAGCTAGAACGG-3’
2. pII-BsrGI-rev: 5’-AGATGAACGGTGTACAGACCAGG-3’
3. pII-ile40-rev: 5’-TAAATTAATGCCGGAGAGGGTAGC-3’
4. pII-SbfI-for: 5’-GACTCCTGCAGGTCTCGGG-3’
5. pII-opdel-for: 5’-ACAATCTTCCTTGATTATCAACCGGGGTACAT-3’
6. pII-opdel-rev: 5’-GGTTGATAATCAAGGAAGATTGTATAAGCAAATATTTAAATTGT-3’
7. pBAD-gIII-XbaI-for: 5’-CGCGGGCTATCTAGAGTGAAAAAATTATTATTCGCAAT-3’
8. pBAD-gIII-BamHI-rev: 5’-CAGGGCCCGGATCCTTAAGACTCCTTATTACGCAG-3’
9. Cys(NNK)_5_ Cys-for: 5’-GCTTCCATGGCCTGCNNKNNKNNKNNKNNKTGCGCGGCGAAAGCGGCC-3’
10. Cys(NNK)_5_ Cys-rev: 5’-GCTCCATGGCCGGCTGGGCCGC-3’

#### Plasmid Construction

All novel plasmid maps are described in the supporting information (Figure S9).

##### CMa13d3

The plasmid CM13d3 (Antibody Design Labs) was digested with SacI (New England Biolabs) for 10 hours at 37 °C. The digestion mixture was then gel-purified and the CMaPylRS/PylT gene cassette (gBlocks, IDT) was ligated into the digested vector using T4 ligase (New England Biolabs). The ligation product was transformed into to Top10 chemically competent cells, then confirmed by Sanger Sequencing.

##### CMa13ile40

The met40ile mutation was generated in *CMa13d3* by performing an overlap extension PCR of the pII region. First, regions were amplified using primer pairs pII-ile40-for/pII-BsrGI-rev and pII-ile40-rev/pII-SbfI-for. The PCR products were then gel extracted, combined, and amplified using primers pII-SbfI-for and pII-BsrGI-rev. The product was then digested for 1 hour at 37 °C with SbfI and BsrGI and ligated into *CMa13d3* that had also been digested with BsrGI and SbfI. The ligation product was transformed into Top10 and confirmed by Sanger sequencing.

##### CMa13opdel

The pII operator deletion in *CMa13d3* was generated by site directed mutagenesis using primers pII-opdel-for and pII-opdel-rev. The PCR product was purified using a midiprep kit (Epoch Biosciences) and then directly transformed into Top10 cells. The deletion was confirmed using Sanger sequencing.

##### pBAD-gIII

A gIII gene fragment containing EcoRI and XbaI restriction cut sites was PCR-amplified from CM13 plasmid (Antibody Design Labs) using primers pBAD-gIII-XbaI-for and pBAD-gIII-BamHI-rev. The gene fragment and pBAD vector were digested with EcoRI and XbaI for 10 hours at 37 °C before being heat-inactivated at 75 °C for 10 min. The expected bands were gel extracted, ligated with T4 ligase for 10 h, and then heat-inactivated at 75 °C for 10 min. The ligation products were then transformed into Top10 chemically competent cells and clones were confirmed by Sanger sequencing.

##### pBAD-sfGFP-134TAG

The *pBAD-sfGFP-134TAG* plasmid has been previously reported.^34^

##### pADL-10b-12mer

The *pADL-10b-12mer* plasmid has been previously reported.^35^

##### pADL-10b-5mer

We constructed this library by undergoing PCR to directly amplify the pADL-10b plasmid using primers Cys(NNK)_5_ Cys-for and Cys(NNK)_5_ Cys-rev. The phagemid template DNA was digested using DpnI at 37°C for 1h. PCR product was digested using NcoI restriction enzyme and ligated using T4 DNA ligase at 16 °C for overnight. The ligation products were heated at 75 °C for 10 min to denature the T4 enzyme and then transformed into Top10F’ electrically competent cells.

##### pET28a-SUMO-ZNRF3-Avi

A gene fragment encoding the extracellular domain of ZNRF3 (residues 56-219) was ordered from IDT and the full sequence can be found in the Supporting Information. This was digested using BamHI and XhoI, then ligated using T4 ligase into a pET28a vector that contained a SUMO-tag upstream of the BamHI site. The plasmid was transformed into Top10 cells and verified with Sanger sequencing.

### 2. Biological Methods

#### Helper Phage Expression

Helper phage plasmids were transformed into Top10 containing *pBAD-gIII*. Cells were then used to inoculate 2xYT media containing antibiotics (100 µg/mL ampicillin, 25 µg/mL Kanamycin) and grew until OD_600_ = 0.4. Phage expression was then induced by addition of 0.2% arabinose and phages expressed for 20 hours at 30 °C. Phages were purified and concentrated using PEG/NaCl precipitations before being titered via the colony-forming unit assay.

#### Phage Quantification

For all experiments, phages were quantified via a colony forming unit assay. In this assay, serial dilutions of the phage solution were prepared in 2xYT media and 10 µL of each dilution was added to 90 µL of log-phase *E. coli* ER2738. Following addition of the phage dilutions, the culture was incubated at 37 °C for 45 min and then 10 µL was spotted in triplicate onto agar selection plates containing 100 µg/mL ampicillin for pADL libraries or 25 µg/mL kanamycin for helper phage and 10 µg/mL tetracycline, which were incubated at 37 °C overnight. The following day, colonies in each spot were counted and this number was used to calculate the number of colony-forming units in the solution.

#### Packaging Selectivity Assay

Helper phage plasmids (*CMa13ile40* or *CM13d3*) were transformed into Top10 containing *pADL-10b-AAKAA*. Cells were then used to inoculate 2xYT media containing antibiotics (100 µg/mL ampicillin, 25 µg/mL Kanamycin) and grew until OD_600_ = 0.5. Phage expression was then induced by addition of 1 mM IPTG and phages expressed for 20 hours at 30 °C. Phages were purified and concentrated using PEG/NaCl precipitations before being titered via the colony-forming unit assay. Packaging of the pADL-AAKAA or helper phage library was detected by formation of ampicillin or kanamycin resistant colonies, respectively.

#### GFP Production with CMa13ile40

The plasmid *pBAD-134TAGsfGFP* was transformed into Top10F’ cells. Cells were then grown to OD_600_ = 0.4 in 2xYT media containing ampicillin (100 µg/mL) and tetracycline (10 µg/mL) before being transfected with CMa13ile40 (MOI = 1) for 45 min at 37 °C. Cells were then pelleted (3750 xg, 15 min) and resuspended in 2xYT supplemented with appropriate antibiotics (100 µg/mL ampicillin, 25 µg/mL kanamycin, 10 µg/mL tetracycline), 0.2% arabinose, 1 mM IPTG, and with or without ncAAs (1 mM). The cells expressed for 16 hours at 37 °C. Following expression, the OD_600_ was measured for each condition. The cells were then pelleted (3750 xg, 15 min) and resuspended in 1 mL of lysis buffer (50 mM NaCl, 50 mM Tris, pH 8.0). Cells were then lysed by three freeze/thaw cycles using liquid N_2_ and a 42 °C heat bath, with vortexing between cycles. The lysate was then clarified by centrifugation (3750 xg, 15 min) and the supernatant was removed to measure GFP fluorescence. Fluorescence was measured using a Neo2 plate reader (485/528 nm excitation/emission) and values were normalized to OD_600_ by dividing the total fluorescence by the measured OD_600_.

#### Amber Enrichment using CMa13ile40

Plasmids encoding the peptide libraries were transformed into electrocompetent Top10F’, reaching an efficiency of 10^10^ total transformants. Library stocks were then inoculated into 1 L of 2xYT supplemented with 100 µg/mL ampicillin and 10 µg/mL tetracycline. The culture grew to OD_600_ = 0.25, then a 5 mL aliquot was removed as a negative control. To the remaining culture, pIII production was induced by addition of 0.2 mM IPTG and incubation for 30 min at 37 °C. Following incubation, 10^10^ total cells were removed from the culture and were infected with CMa13ile40 (MOI: 1) for 45 min at 37 °C. Cells were then pelleted (3750 xg, 15 min) and resuspended in 2xYT containing tetracycline (10 µg/mL), ampicillin (100 µg/mL), kanamycin (25 µg/mL), 1 mM IPTG, and 5 mM of ncAA (BocK or AllocK). The cells were diluted 10-fold in the final expression media to allow for selection of cells that were kanamycin resistant. Control expressions were performed in 25 mL final volume of the expression media with or without ncAA depending on the control. Phages were expressed at 30 °C for 20 hours, then the cells were pelleted (3750 xg, 15 min) and phages were concentrated from the supernatant through two subsequent PEG/NaCl precipitations. Ultimately, phages were resuspended in 1 mL of binding buffer (10 mM HEPES, 150 mM NaCl, 10 mM MgCl_2_, 1 mM KCl, pH 7.4). Residual bacteria were killed by incubation at 65 °C for 15 min and phages were titered in ER2738 using the colony-forming unit assay. Phage yields from the controls were normalized to total expression volumes of the ncAA containing solutions.

#### Illumina Sequencing of Phage Libraries

Phage libraries were isolated by infection into ER2738 and growth of cells overnight in 2xYT containing ampicillin (100 µg/mL) and tetracycline (10 µg/mL). Following infection, plasmid DNA was isolated using a miniprep DNA extraction column. A four step PCR cycle was used to amplify the library region out of the original phagemid library using primers NGS-F1 and NGS-R1, as has been previously described. The amplicons were purified and extracted from a 3% agarose gel according to a GenCatch gel extraction kit, then indices were attached using a subsequent PCR with NGS-i7 and NGS-i5 primers. To identify the rounds of selection, each round contained a unique combination of i7 or i5 indices. The PCR products were purified using a GenCatch gel extraction kit and submitted to the Genomics and Bioinformatics center at Texas A&M University for sequencing on an Illumina MiSeq (1M Reads, 2x150bp). Sequences were analyzed for enrichment in R, and all scripts are available in the Supporting Information.

#### Expression of ZNRF3

Plasmid *pET28a-SUMO-ZNRF3-Avi* was cotransformed into *E. coli* (BL21) along with a plasmid expressing BirA. The cells were then inoculated into 2xYT containing 50 µg/mL kanamycin and 20 µg/mL chloramphenicol and grew at 37 °C until OD = 0.5. Expression was then induced by addition of 1 mM IPTG and 50 µM biotin, then the protein was for 18 h at 16 °C. The cells were then pelleted (3750 x *g*, 15 min) and then resuspended in buffer A (50 mM Tris-HCl, 10% glycerol, 150 mM NaCl, 25 mM imidazole, pH 7.5, 20 mL/liter of expression). The cells were then lysed by sonication (1 s on, 1 s off, 5 min, 60% A, repeated once) and the lysate was clarified by centrifugation (25000 x *g*, 20 min). The supernatant was then incubated with a Ni-NTA column for 45 min at 4 °C. The column was then washed with 5 CV of buffer A before being eluted using 2 CV of buffer B (buffer A + 240 mM imidazole). The elution was then concentrated using an Amicon centrifugal filter before being desalted into buffer A on a HiPrep 26/10 column. The SUMO tag was then cleaved by addition of 30 µg/mL SUMO protease and incubation for 48 h at 4 °C. The protein was then loaded onto a second Ni-NTA column for 45 min at 4 °C, and the flowthrough was collected and concentrated to 1 mL by centrifugation. Pure protein was then isolated using size exclusion chromatography (HiLoad 16/600, Superdex 75 pg) in buffer C (50 mM Tris-HCl, pH 7.5, 10% (v/v) glycerol, 150 mM NaCl, 0.5 mM EDTA). Pure protein was stored in storage buffer (50 mM Tris-HCl, pH 7.5, 50% (v/v) glycerol, 150 mM NaCl) at -80 °C before being used for assays.

#### Affinity Selection Against ZNRF3

##### Phage Selection

Streptavidin coated magnetic beads (100 µL, 50% slurry, Cytiva Sera-Mag) were transferred to a 1.5 mL tube, washed three times with 1 mL of Binding Buffer (10 mM HEPES, 150 mM NaCl, 10 mM MgCl_2_, 1 mM KCl, pH 7.4), resuspended in 100 uL of Binding Buffer, and split into two tubes. 500 pmol of biotinylated ZNRF3 was added to 1 mL of Binding Buffer in one of the tubes and an equal volume of Binding Buffer was added to the other tube (hereafter referred to as + tube and – tube, respectively). The beads/protein mixture was incubated with rocking for 15 min at room temperature. The supernatant was then removed, and the beads were washed with Binding Buffer (3 × 1 mL). 0.25 mL of 5x blocking buffer (Binding Buffer + 5% BSA + 0.5% Tween 20) was added to each tube and phage solution, and they incubated at room temperature with end-over-end rotation for 30 min. The blocking buffer was removed from the – tube, and the purified phage library was incubated with the beads for 30 min at room temperature as a negative selection. The supernatant was then transferred to the resin of the + tube and left to incubate at room temperature for 30 min. After 30 min the supernatant was removed, and the resin was washed with Wash Buffer (Binding Buffer + 0.1% Tween 20) to remove nonspecifically bound phages. For the first round of selection, 3 × 1 mL washes were performed, then 5 × 1 mL and 10 × 1 mL washes were done to increase stringency in the second and third rounds, respectively. During each washing step the resin was completely resuspended by pipetting up and down. To remove phages binding to the polypropylene tube, the resin was transferred to fresh tubes after every other wash. After the last wash, phages were eluted by incubating for 15 min with 100 µL of Elution Buffer (50 mM glycine, pH 2.2). The supernatant was then removed and immediately added to 50 µL of Neutralization Buffer (1 M Tris, pH 8.0). The neutralized elution was immediately used for amplification.

#### Phage Amplification/Expression

A small aliquot (10 µL) of the phage elution was removed for quantification of phages. The remaining solution was added to an actively growing culture of *E. coli* Top10F’ (OD_600_ = 0.5-0.6) in 20 mL of 2xYT containing 10 µg/mL tetracycline for 45 min at 37 °C with rotation. After 45 min the cells were pelleted (3750 × *g*, 15 min), resuspended in 200 mL of 2xYT containing 100 µg/mL ampicillin and 10 µg/mL tetracycline, and amplified overnight at 37 °C. The following day, phagemids were extracted from the amplified culture using a commercial plasmid purification kit. The remaining culture was then inoculated into 1 L of fresh 2xYT containing 100 µg/mL ampicillin and 10 µg/mL tetracycline and grown to OD_600_ = 0.4. Cells to cover library size of the eluted phages were removed (typically 10^9^ total cells, 20 mL culture) and infected with CMa13ile40 (MOI: 1.2) for 45 min at 37 °C. The cells were then pelleted (3750 × *g*, 15 min) and resuspended in 200 mL of 2xYT containing 100 µg/mL ampicillin, 10 µg/mL tetracycline, 25 µg/mL kanamycin, l mM IPTG, and 5 mM of ncAA (BocK or AllocK). As a negative control, a 20 mL expression lacking the appropriate ncAA was also performed. Phages were then expressed for 20 hours at 30 °C, before being purified and concentrated by PEG/NaCl precipitations. Ultimately, phages were resuspended in 1 mL of binding buffer and titered using the colony-forming unit assay.

#### Biolayer Interferometry Assays

Biolayer interferometery (BLI) experiments were performed using streptavidin coated biosensors (Sartorius Bio) on an Octet Red 96 biolayer interferometer (Sartorius Bio). All experiments were performed in assay buffer (10 mM HEPES, 150 mM NaCl, 0.01% Tween 20, pH 7.5). Biotinylated ZNRF3 (28 µg/mL in assay buffer) was loaded onto the sensor to 4 nm of loading over 10 min, then the sensors were quenched for one minute with a 1% BSA solution in assay buffer. Peptides were serially diluted in assay buffer containing 0.8% DMF and the association and dissociation to ZNRF3 were measured over 3 and 5 min, respectively. Between each experiment, sensors were regenerated using 10 mM glycine, pH 1.5. Binding curves were double-referenced against a protein-only sensor and a parallel unloaded sensor to correct for any nonspecific binding to the sensors. All curves were fit in the Octet Red Kinetic Analysis program using a 1:1 protein:ligand binding model.

### 3. Synthetic Methods

#### Peptide Synthesis

##### Coupling Conditions

Initial sequences of peptides were synthesized on a low loading ProTide rink amide resin (CEM #R002) using a MultiPep 2 Peptide Synthesizer (CEM). All amino acid derivatives were standard derivatives for Fmoc peptide synthesis commercially purchased from Chem-Impex. For cysteine oxidation, Fmoc-Cys(mmt)-OH was used as the derivative. Fmoc-amino acids were deprotected using 20% piperidine in DMF (2 × 2 mL, 5 min at r.t.). Amino acids were coupled using HATU/NMM double coupling cycles (4.2 eq amino acid, 4.2 eq HATU, 8 eq NMM, 30 min r.t.).

##### Cysteine Oxidation

To deprotect the Cys(mmt) on-resin, peptides were washed (6 × 3 mL) with 94:1:5 DCM:TFA:TIPS. An orange color in the flowthrough indicated loss of the methoxytrityl protecting group. The free cysteines were then oxidized using N-chlorosuccinimide, as previously reported (1.4 mg/mL, 1.05 eq, 15 min). Resins were then washed with DMF, followed by DCM and dried in vacuo.

##### Peptide Cleavage

The peptides were cleaved from resin by agitating for 3 h in 3 mL of 92.5:2.5:2.5 TFA:H2O:TIS. The products were then filtered, and peptides were precipitated out of the filtrate using cold ether. The precipitates were collected by centrifugation and washed again with cold ether, then lyophilized.

##### Boc Protection of Lysine

For peptides ZPB1 and ZPB2, peptides were cleaved with the N-terminus Fmoc-protected. The free lysine was then protected by incubation with boc-anhydride (3 eq for ZPB1, 1 eq for ZPB2) and NMM (4 eq for ZPB1, 1.3 eq for ZPB2) for 24 hours at room temperature. The N-terminus was then deprotected by incubation with 20% piperidine in DMF (2 × 500 µL, 5 min). Peptide solutions were then diluted with water and lyophilized.

##### Purification of Peptides

Crude peptides were dissolved in 1:2:2 DMSO:ACN:H_2_O and then purified using an acetonitrile:water gradient on a reverse-phase preparative HPLC column (Shim-pack GIS C18 column, 20 × 250 mm, 10 µm, Shimadzu). Purified peptides were analyzed for purity using high resolution mass spectrometry. All LC-MS spectra can be found in the Supporting Information (Figures S10-S16).

## ACKNOWLEDGMENTS

This work was support in part by National Institutes of Health (Grants R35GM145351 to W.R.L.), Welch Foundation (Grant A-1715 to W.R.L.), the Texas A&M X-Grants Mechanism, and the Texas A&M Chancellor EDGES Fellowship.

