## Supporting Information for "An Amber-Encoding Helper Phage for More Efficient Phage Display of Noncanonical Amino Acids"

### SUPPLEMENTARY FIGURES

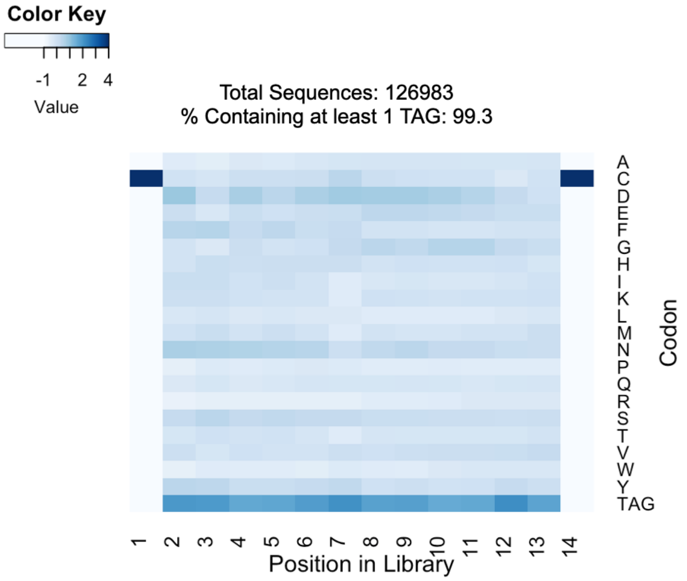


**Figure S1:** Heat map for the 12mer amber enriched library. Enrichment values were calculated by (%library-%expected)/%expected.

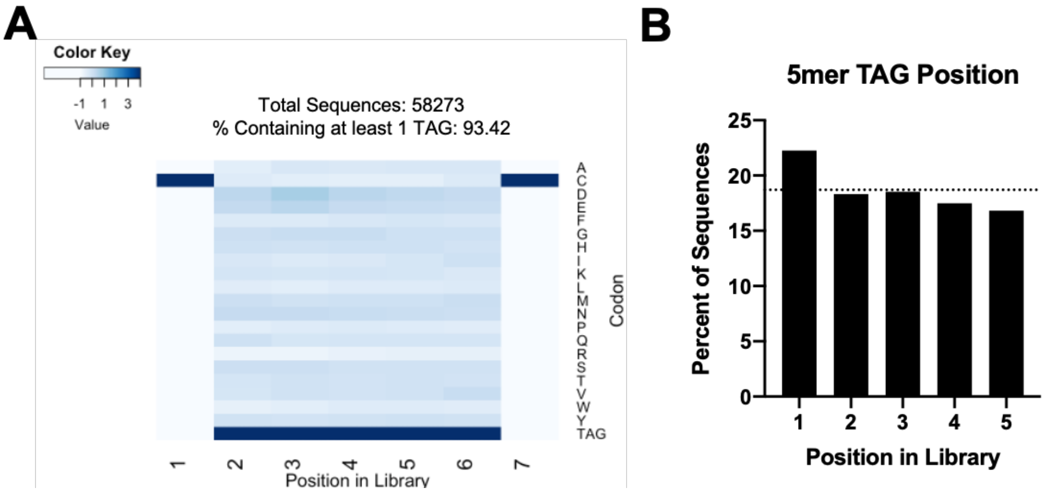


**Figure S2:** A) Heat map for the 5mer amber enriched library. Enrichment values were calculated by (%library-%expected)/%expected. B) % of sequences containing TAG codons at each of the 5 positions in the library. The dashed line corresponds to the expected values of a truly randomized library with the observed amber percentage.

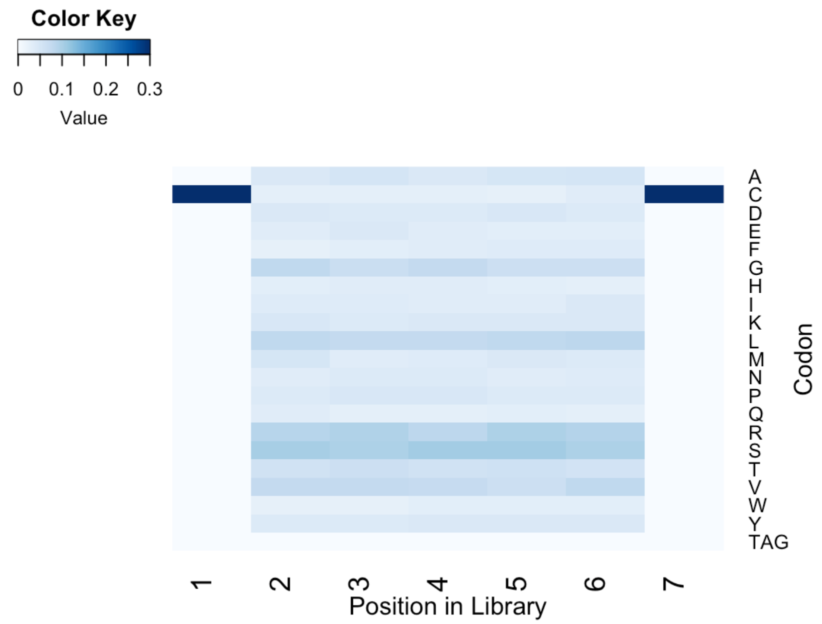


**Figure S3:** Heat map for sequences of the 5mer amber enriched library that do not contain TAG. Enrichment values were calculated by (%library-%expected)/%expected.

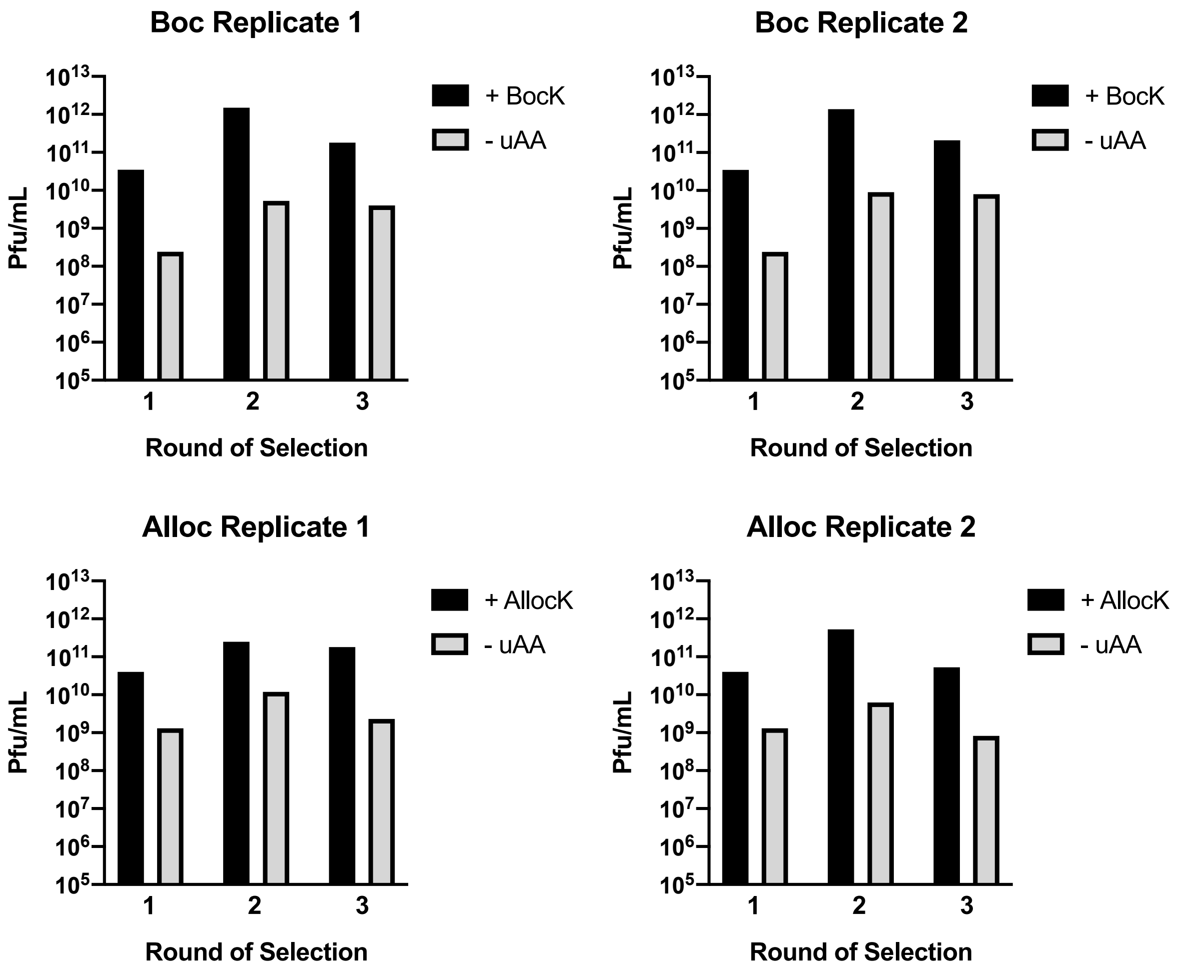


**Figure S4:** Phage expression titers for each replicate for selections against ZNRF3 using BocK and AllocK. Each phage yield is normalized to a 200 mL final expression volume.

**A**

A1R3

| | V1 | V2 | V3 | V4 | V5 | V6 | V7 | V8 | V9 | V10 | V11 | V12 | V13 | V14 | n | percent |
|---|---|---|---|---|---|---|---|---|---|---|---|---|---|---|---|---|
| ZPA1 | C | H | M | TAG | E | Y | D | N | V | L | W | E | V | C | 74323 | 86.8533299054608 |
| | C | I | G | S | I | V | I | TAG | D | G | Y | T | W | C | 9867 | 11.5305061175838 |
| | C | D | S | L | Y | TAG | D | S | F | G | Y | W | W | C | 721 | 0.842555478947799 |
| | C | Q | I | V | E | TAG | H | G | Y | M | W | A | Y | C | 43 | 0.0502494945835719 |
| | C | E | S | V | Q | TAG | L | D | Y | W | W | Y | Y | C | 34 | 0.0397321585079406 |
| | C | H | M | TAG | E | Y | D | N | V | L | C | E | V | C | 20 | 0.0233718579458474 |
| | C | H | M | TAG | G | Y | D | N | V | L | W | E | V | C | 20 | 0.0233718579458474 |
| | C | H | M | TAG | E | C | D | N | V | L | W | E | V | C | 18 | 0.0210346721512627 |
| | C | H | M | TAG | E | Y | D | N | V | S | W | E | V | C | 18 | 0.0210346721512627 |
| | Y | H | M | TAG | E | Y | D | N | V | L | W | E | V | C | 18 | 0.0210346721512627 |
| | C | H | I | TAG | E | Y | D | N | V | L | W | E | V | C | 17 | 0.0198660792539703 |
| | C | H | M | TAG | E | Y | D | N | V | L | R | E | V | C | 15 | 0.0175288934593856 |
| | C | H | M | TAG | E | Y | D | N | V | L | W | E | L | C | 13 | 0.0151917076648008 |
| | C | H | M | TAG | E | Y | V | N | V | L | W | E | V | C | 13 | 0.0151917076648008 |
| | C | H | M | Y | E | Y | D | N | V | L | W | E | V | C | 13 | 0.0151917076648008 |

**B**

A2R3

| | V1 | V2 | V3 | V4 | V5 | V6 | V7 | V8 | V9 | V10 | V11 | V12 | V13 | V14 | n | percent |
|---|---|---|---|---|---|---|---|---|---|---|---|---|---|---|---|---|
| ZPA1 | C | H | M | TAG | E | Y | D | N | V | L | W | E | V | C | 88368 | 91.611030478955 |
| | C | I | G | S | I | V | I | TAG | D | G | Y | T | W | C | 7043 | 7.30147211279287 |
| | C | D | S | L | Y | TAG | D | S | F | G | Y | W | W | C | 428 | 0.443707236160066 |
| | Y | H | M | TAG | E | Y | D | N | V | L | W | E | V | C | 27 | 0.0279908770474808 |
| | C | E | S | V | Q | TAG | L | D | Y | W | W | Y | Y | C | 19 | 0.0196972838482272 |
| | C | H | M | TAG | E | Y | D | N | V | L | C | E | V | C | 18 | 0.0186605846983205 |
| | C | H | M | TAG | E | Y | D | N | V | L | R | E | V | C | 18 | 0.0186605846983205 |
| | C | H | M | TAG | E | Y | D | N | V | S | W | E | V | C | 16 | 0.0165871863985072 |
| | C | T | I | I | Y | TAG | E | G | V | K | W | V | K | C | 16 | 0.0165871863985072 |
| | R | H | M | TAG | E | Y | D | N | V | L | W | E | V | C | 14 | 0.0145137880986938 |
| | C | H | M | TAG | E | Y | N | N | V | L | W | E | V | C | 13 | 0.0134770889487871 |
| | C | H | M | TAG | E | C | D | N | V | L | W | E | V | C | 12 | 0.0124403897988804 |
| | C | H | M | TAG | E | Y | D | N | V | L | TAG | E | V | C | 12 | 0.0124403897988804 |
| | C | H | I | TAG | E | Y | D | N | V | L | W | E | V | C | 11 | 0.0114036906489737 |
| | C | H | M | TAG | E | Y | D | D | V | L | W | E | V | C | 11 | 0.0114036906489737 |

**Figure S5:** Next-generation sequencing data for the selection against ZNRF3 using AllocK. Sequences from the third round of selection for replicates 1 and 2 are shown in (A) and (B) respectively. TAG codons correspond to AllocK incorporation.

**A**

B1R3

| | V1 | V2 | V3 | V4 | V5 | V6 | V7 | V8 | V9 | V10 | V11 | V12 | V13 | V14 | n | percent |
|---|---|---|---|---|---|---|---|---|---|---|---|---|---|---|---|---|
| ZPB1 | C | E | V | T | TAG | D | Y | T | W | S | V | W | P | C | 48915 | 50.8250036366659 |
| ZPB2 | C | Y | S | T | S | TAG | H | G | Y | M | W | V | T | C | 26640 | 27.6802227717628 |
| | C | N | Y | V | H | TAG | G | G | Y | D | W | Q | H | C | 6573 | 6.82965856902392 |
| | C | I | G | S | I | V | I | TAG | D | G | Y | T | W | C | 5425 | 5.63683215228279 |
| | C | D | TAG | Y | I | I | S | D | G | G | Y | L | W | C | 3177 | 3.30105359406496 |
| | C | D | S | L | Y | TAG | D | S | F | G | Y | W | W | C | 693 | 0.720059849130317 |
| | C | S | N | I | V | TAG | D | K | W | V | Y | E | M | C | 673 | 0.699278901103468 |
| | C | V | L | TAG | F | D | G | N | Y | S | W | E | V | C | 645 | 0.67018557386588 |
| | C | T | I | E | F | TAG | Y | D | Y | D | W | T | V | C | 610 | 0.633818914818894 |
| | C | A | Y | T | W | TAG | Y | D | Y | E | W | E | F | C | 525 | 0.545499885704786 |
| | C | T | I | I | Y | TAG | E | G | V | K | W | V | K | C | 497 | 0.516406558467197 |
| | C | D | TAG | I | W | R | D | G | Y | W | W | L | A | C | 319 | 0.331456121028241 |
| | C | S | S | S | T | I | S | W | TAG | A | K | L | E | C | 204 | 0.21196566987386 |
| | C | E | S | V | Q | TAG | L | D | Y | W | W | Y | Y | C | 127 | 0.131959019970491 |
| | C | Q | G | R | Y | TAG | F | Y | D | G | Y | L | W | C | 104 | 0.108060929739615 |

**B**

B2R3

| | V1 | V2 | V3 | V4 | V5 | V6 | V7 | V8 | V9 | V10 | V11 | V12 | V13 | V14 | n | percent |
|---|---|---|---|---|---|---|---|---|---|---|---|---|---|---|---|---|
| | C | N | Y | V | H | TAG | G | G | Y | D | W | Q | H | C | 33026 | 42.8708655693442 |
| ZPB1 | C | E | V | T | TAG | D | Y | T | W | S | V | W | P | C | 22620 | 29.362895269744 |
| ZPB2 | C | Y | S | T | S | TAG | H | G | Y | M | W | V | T | C | 9383 | 12.1800197310348 |
| | C | T | I | E | F | TAG | Y | D | Y | D | W | T | V | C | 5099 | 6.61898333246794 |
| | C | V | L | TAG | F | D | G | N | Y | S | W | E | V | C | 2497 | 3.24134171036918 |
| | C | I | G | S | I | V | I | TAG | D | G | Y | T | W | C | 1839 | 2.3871955968638 |
| | C | A | Y | T | W | TAG | Y | D | Y | E | W | E | F | C | 555 | 0.720442390570642 |
| | C | H | T | I | M | TAG | D | G | Y | A | W | V | R | C | 342 | 0.443948283919207 |
| | C | S | N | I | V | TAG | D | K | W | V | Y | E | M | C | 219 | 0.284282673035983 |
| | C | E | F | I | M | TAG | Q | K | Y | A | W | E | K | C | 203 | 0.26351316267719 |
| | C | M | T | A | M | TAG | W | G | Y | E | W | T | H | C | 134 | 0.173944649254894 |
| | C | V | TAG | S | H | L | S | Y | E | W | Q | F | D | C | 97 | 0.125915156550184 |
| | C | E | S | V | Q | TAG | L | D | Y | W | W | Y | Y | C | 87 | 0.112934212575939 |
| | C | E | V | T | TAG | D | Y | T | W | S | V | W | S | C | 38 | 0.0493275871021341 |
| | C | D | P | V | W | TAG | D | G | V | S | W | L | M | C | 34 | 0.0441352095124357 |

**Figure S6:** Next-generation sequencing data for the selection against ZNRF3 using BocK. Sequences from the third round of selection for replicates 1 and 2 are shown in (A) and (B) respectively. TAG codons correspond to BocK incorporation.

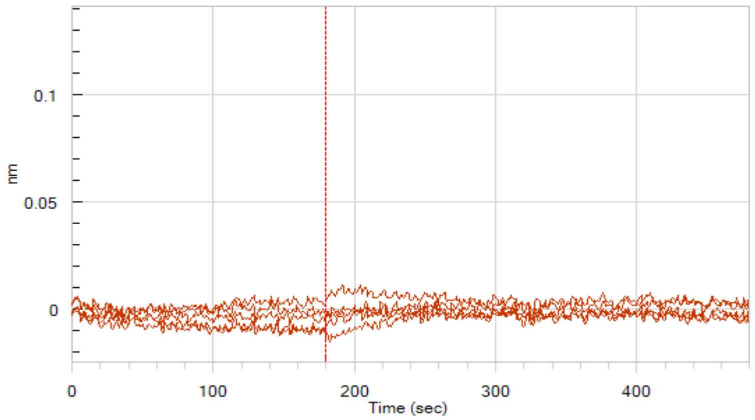


**Figure S7:** BLI data for the binding of ZPA1Abu to ZNRF3. Concentrations of peptide tested: 40, 20, 10, 5, and 2.5 *μM.*

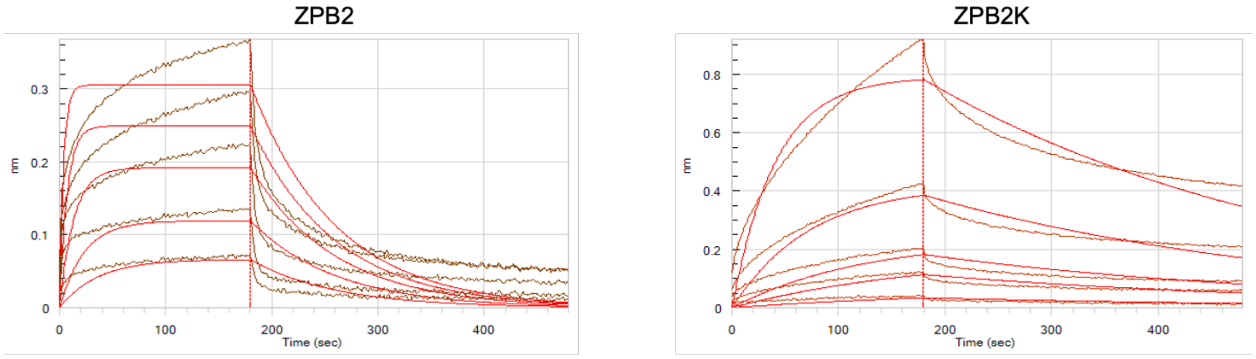


Concentrations (μM): 40, 20, 10, 5, 2.5
$K_D$: 1.5 ± 0.4 μM
N = 3

Concentrations (μM): 40, 20, 10, 5, 2.5
$K_D$: 4.8 ± 0.2 μM
N = 3

**Figure S8:** BLI traces for ZPB2 and ZPB2K binding to ZNRF3. Experimental data are shown in dark red, while fitted curves are shown in light red. $K_D$ values are given as mean ± s.d. of three biological replicates.

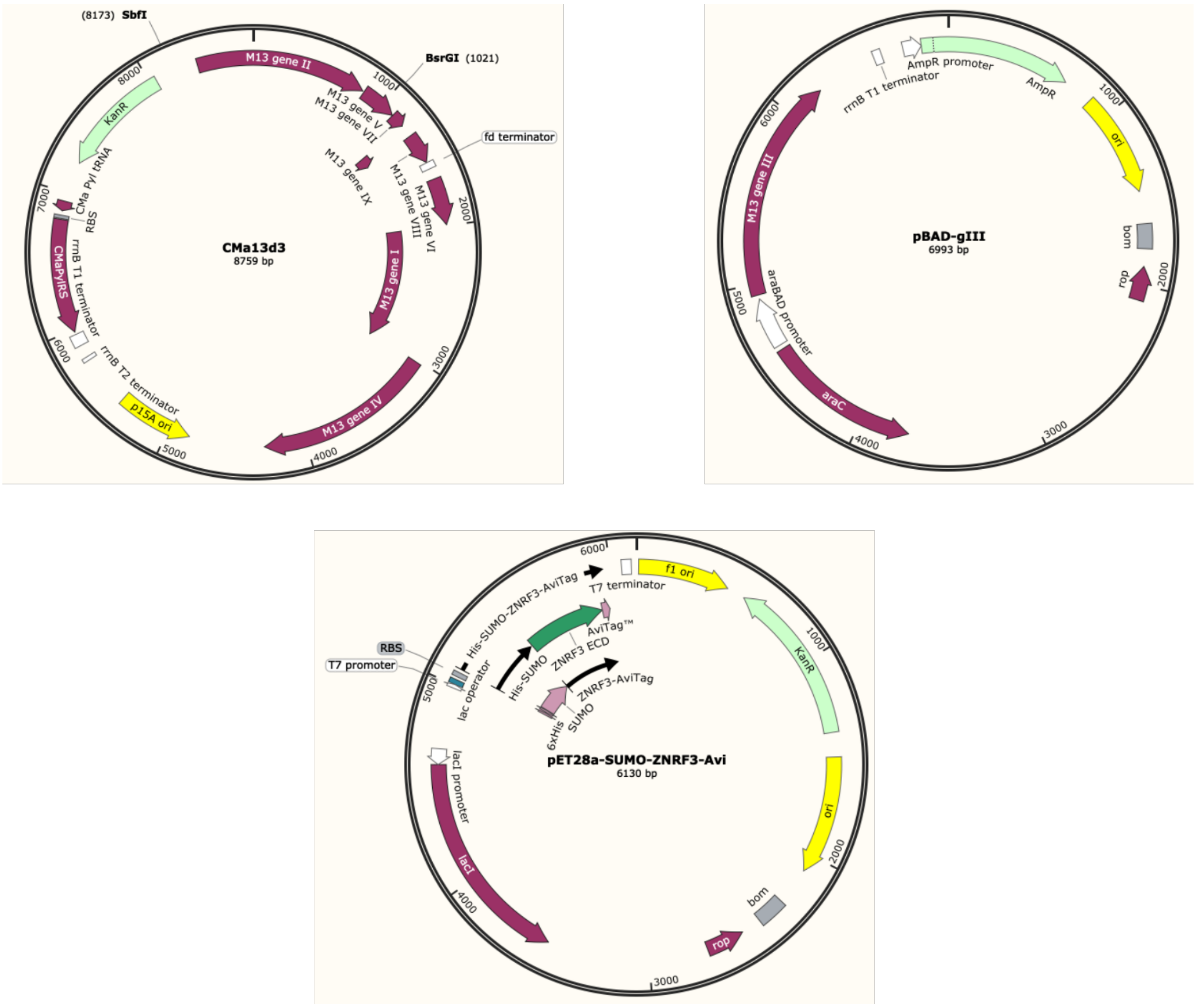


**Figure S9:** Novel plasmid maps for the amber encoding helper phages (CMa13d3), expression of gIII protein (pBAD-gIII) and expression of the extracellular domain of ZNRF3 (pET28a-SUMO-ZNRF3-Avi).

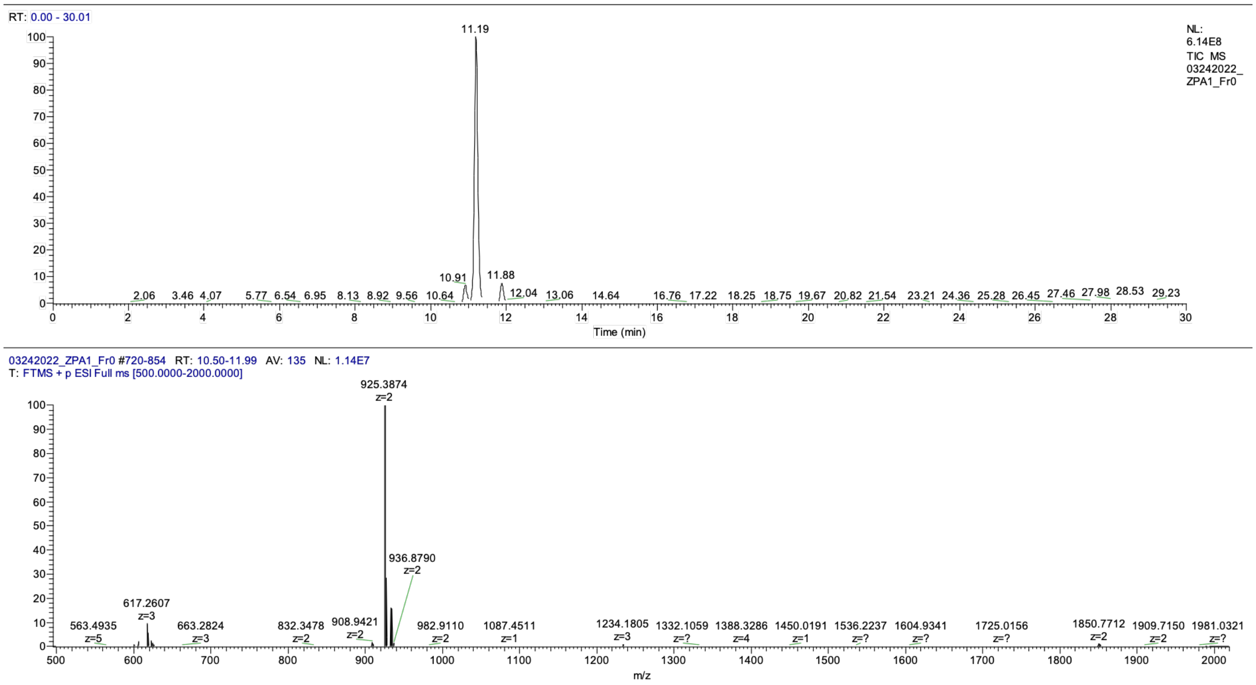


**Figure S10:** LC-MS data for ZPA1. The TIC is shown on top and extracted data for the isolated peak on bottom.

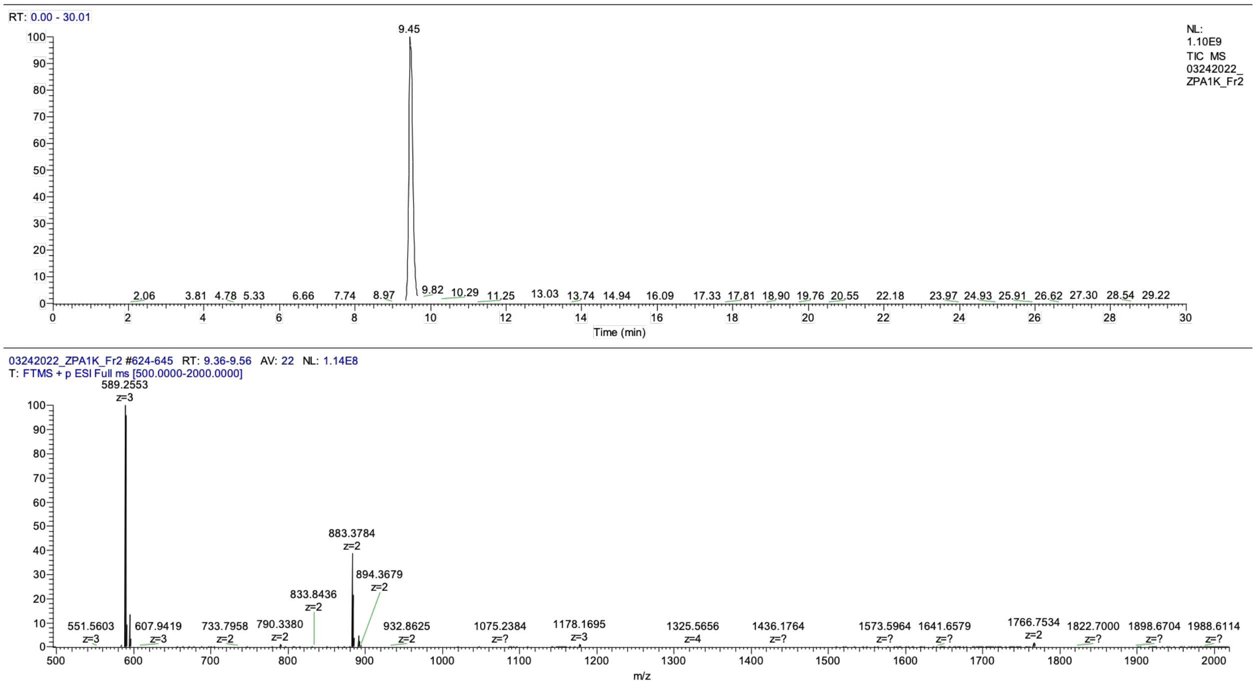


**Figure S11:** LC-MS data for ZPA1K. The TIC is shown on top and extracted data for the isolated peak on bottom.

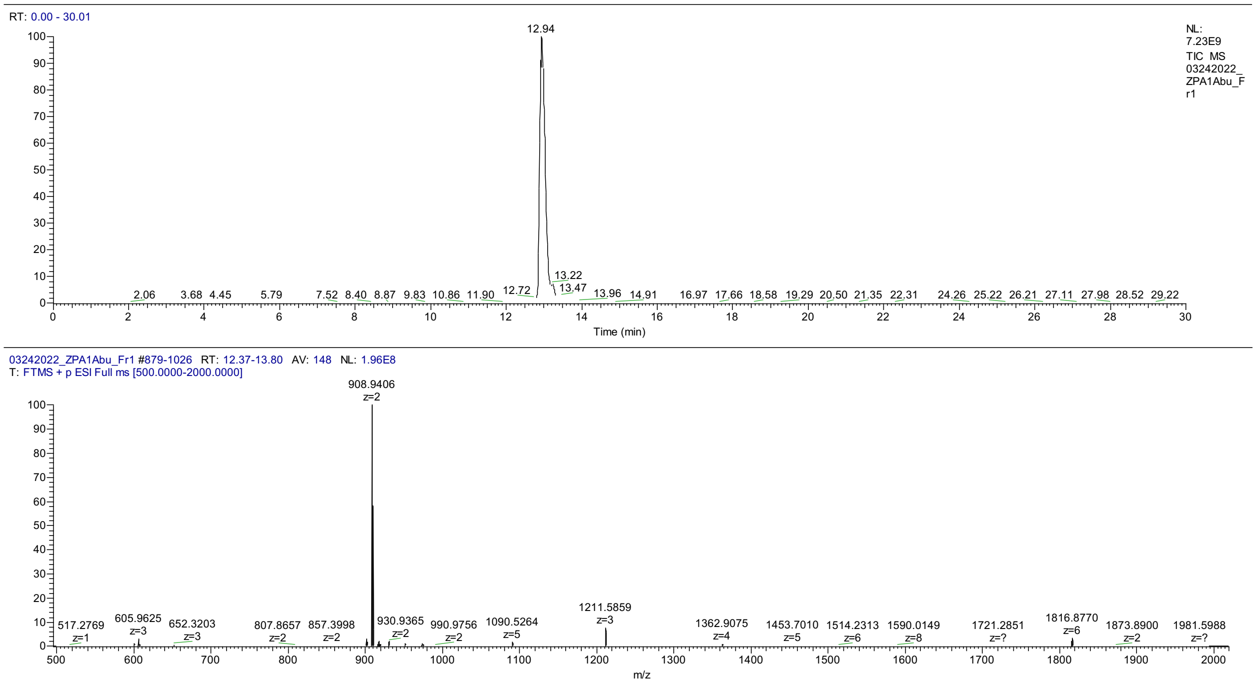


**Figure S12:** LC-MS data for ZPA1Abu. The TIC is shown on top and extracted data for the isolated peak on bottom.

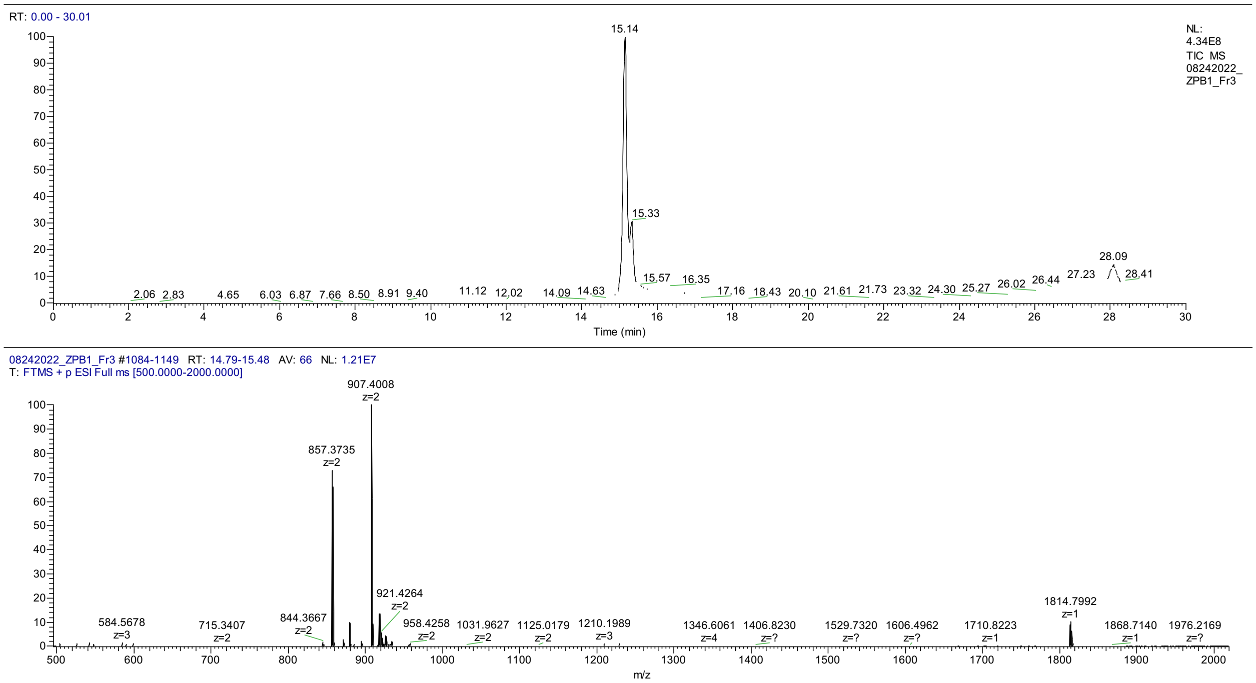


**Figure S13:** LC-MS data for ZPB1. The TIC is shown on top and extracted data for the isolated peak on bottom.

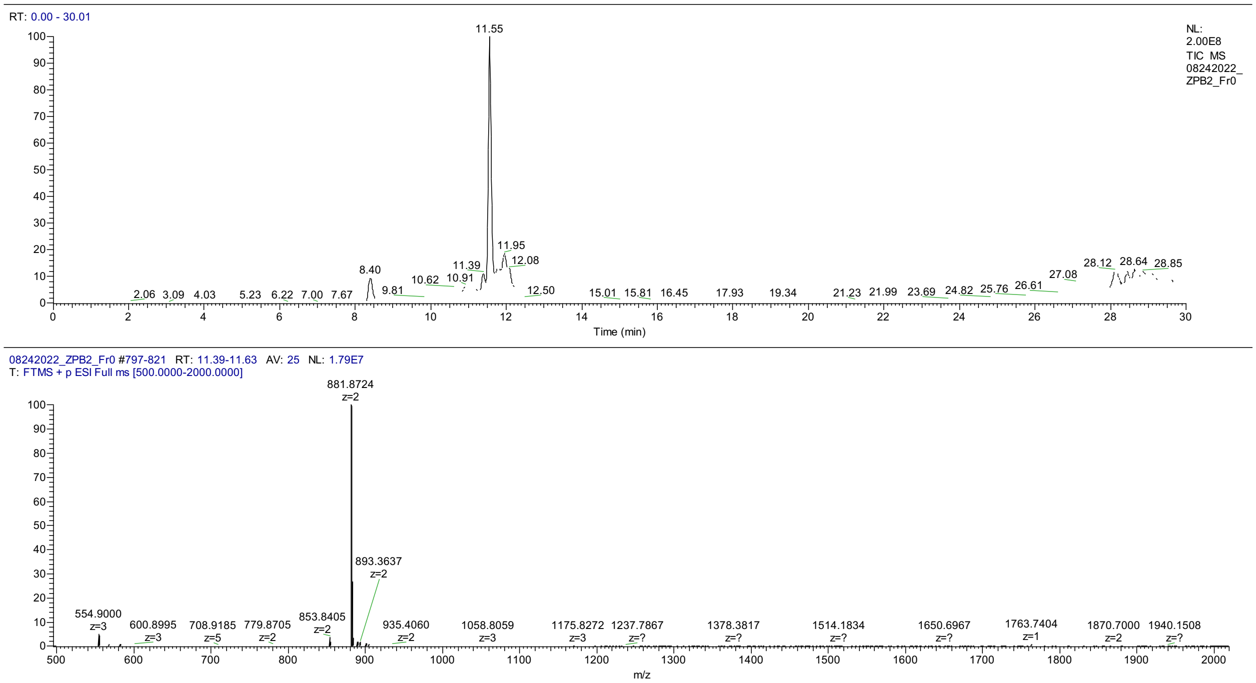


**Figure S14:** LC-MS data for ZPB2. The TIC is shown on top and extracted data for the isolated peak on bottom.

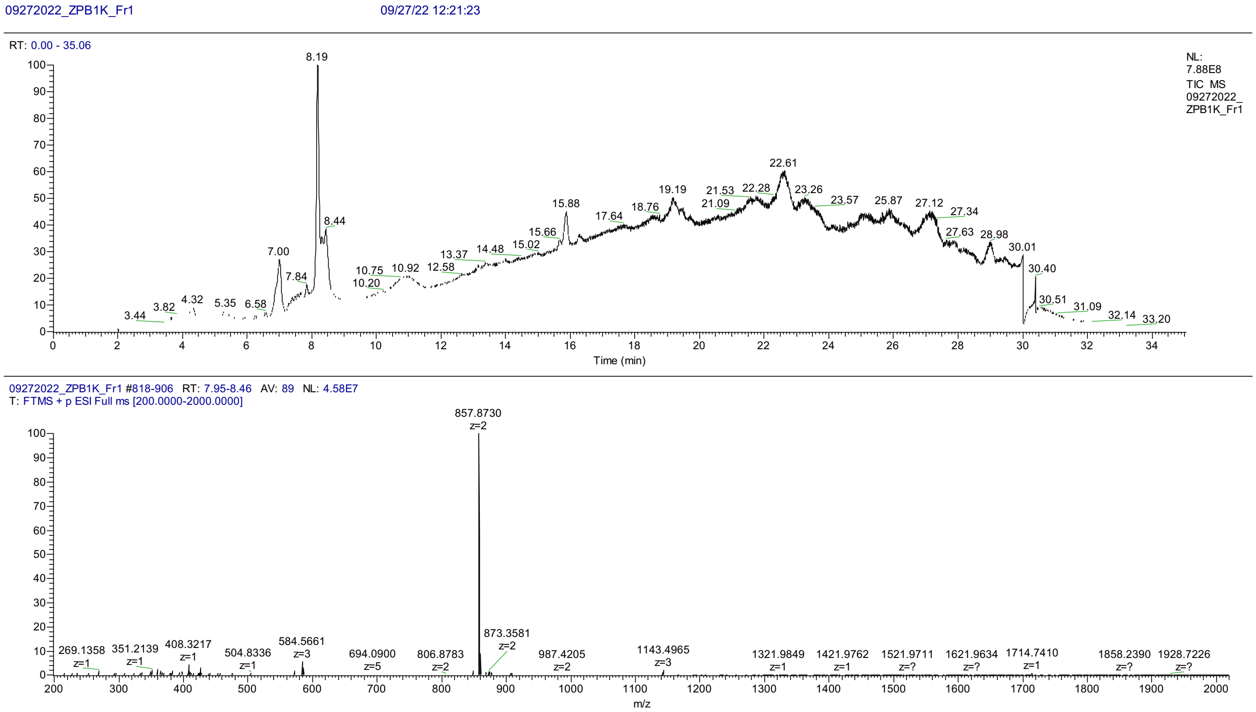


**Figure S15:** LC-MS data for ZPB1K. The TIC is shown on top and extracted data for the isolated peak on bottom.

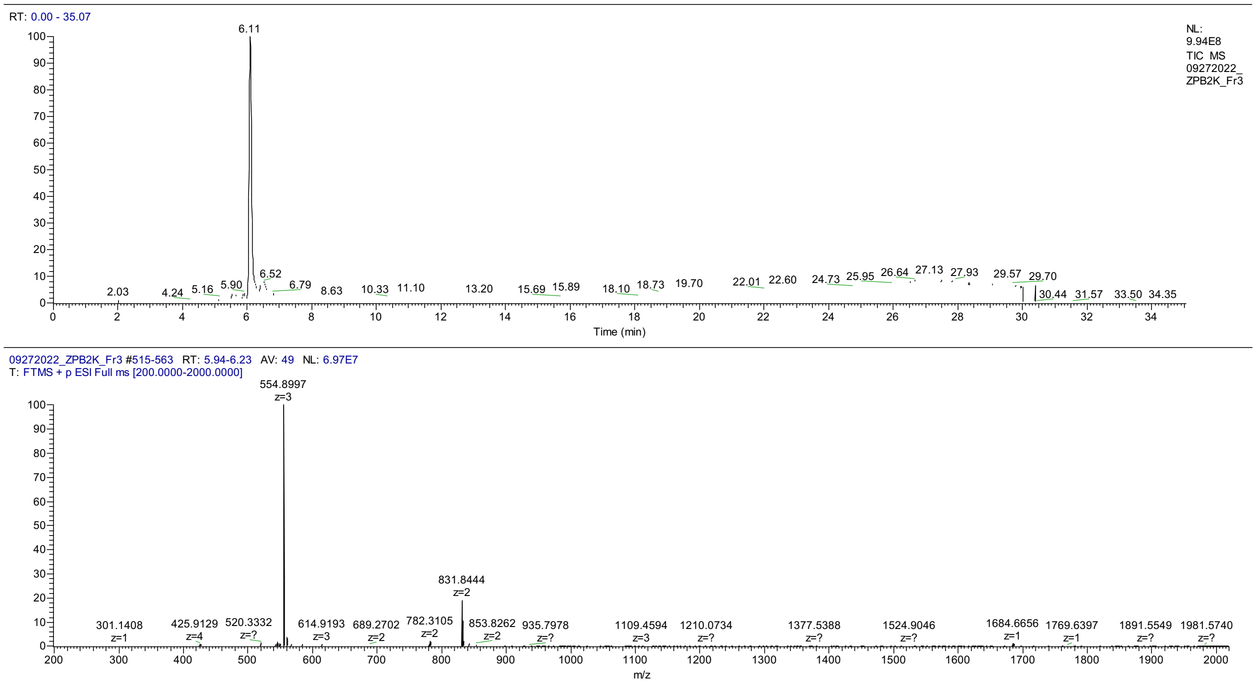


**Figure S16:** LC-MS data for ZPB2K. The TIC is shown on top and extracted data for the isolated peak on bottom.

**SUPPLEMENTARY TABLES**

**Table S1:** High-Resolution Mass Spectrometry Data for Synthesized Peptides

| Peptide | Sequence | Expected Mass (Da) | Observed Mass (Da) |
|---|---|---|---|
| ZPA1 | **C**HMXEYDNVLWEV**C** | 1848.76 | 1848.77 |
| ZPA1K | **C**HMKEYDNVLWEV**C** | 1764.74 | 1764.74 |
| ZPA1Abu | AbuHMXEYDNVLWEVAbu | 1814.87 | 1814.88 |
| ZPB1 | **C**EVTBDYTWSVWP**C** | 1812.79 | 1812.80 |
| ZPB2 | **C**YSTSBHGYMWVT**C** | 1761.73 | 1761.74 |
| ZPB1K | **C**EVTKDYTWSVWP**C** | 1712.73 | 1712.74 |
| ZPB2K | **C**YSTSKHGYMWVT**C** | 1661.68 | 1661.68 |

**C**: Disulfide linked cysteines; X: AllocK; Abu: Aminobutyric acid; B: BocK

**SUPPLEMENTARY SCRIPTS**

**Supplementary Script 1: Amino Acid Analysis**

```
library(microseq)
library(RColorBrewer)
library(dplyr)
library(stringr)
library(gplots)
NNK7Ffilt <-
 readFastq("/Users/traehampton/Documents/Research/Sequencing
 Results/Next Gen Sequencing/22437Wns_N22168/22437Wns_5mer-pp-
 R2_S3_L001_R1_001.fastq")
NNK7Rfilt <-
 readFastq("/Users/traehampton/Documents/Research/Sequencing
 Results/Next Gen Sequencing/22437Wns_N22168/22437Wns_5mer-pp-
 R2_S3_L001_R2_001.fastq")
#Define the following variables
libraryseq <- "GCCCAG.{33}GCGGCG.{6}" #change this regex to match
 specific library
beginning <- 19 #beginning of library in DNA string
lib <- 7 #number of codons in the library region
end <- beginning+lib*3
initialcodon <- beginning%/%3*4-1
endcodon <- initialcodon + lib*4
del <- beginning%/%3 #number of codons before library
aa <-
 c("A","C","D","E","F","G","H","I","K","L","M","N","P","Q","R","S","T",
 "V","W","Y","TAG")

#slices out matches that contain start followed by 24 bases to reverse
 primer
NNK7Ffilt21 <- gregexpr(libraryseq,NNK7Ffilt[[2]],extract = TRUE)
NNK7Rrevcomp <- reverseComplement(NNK7Rfilt[[2]],reverse = TRUE)
 #gives reverse complement of reverse reads
NNK7Rcompfilt21 <- gregexpr(libraryseq,NNK7Rrevcomp,extract = TRUE)

#this compares the forward and reverse strands, only allowing for one
 mismatch in the primers, no mismatches allowed in the library region
n <- length(NNK7Ffilt21)
NNK7Fgood <- vector()
for(i in c(1:n)){
  if(NNK7Ffilt21[[i]] == NNK7Rcompfilt21[[i]]){
    NNK7Fgood[i] <- NNK7Ffilt21[[i]]
  }
  else{
```

```
    split <- strsplit(c(NNK7Ffilt21[[i]],NNK7Rcompfilt21[[i]]), split
 = "")
    diff <- which(split[[1]] != split[[2]])
    if(length(diff) < 2 && length(diff) > 0){
      for(x in c(1:length(diff))){
        if(diff[[x]] < beginning || diff[[x]] > end){
          NNK7Fgood[i] <- NNK7Ffilt21[[i]]
        }
        else{
          NNK7Fgood[i] <- ""
        }
      }
    }
    else{
      NNK7Fgood[i] <- ""
    }
  }
}
NNK7Fgood <- as.data.frame(NNK7Fgood)
NNK7Fgood <- NNK7Fgood[!apply(is.na(NNK7Fgood) | NNK7Fgood == "", 1,
 all),]

#this separates nucleotides into codons
codons <- gsub("(...)", "\\1 \\2", NNK7Fgood)

#this creates dataframe of sequences with reads organized by frequency
seqcount <- as.data.frame(sort(table(codons), decreasing = TRUE))

#this generates a matrix that contains amino acids in library region
l <- length(codons)
AAs <- matrix(0,l,lib)
AA <- gregexpr("\\s(TT[TC])",codons,useBytes = FALSE)
  l <- length(AA)
  for(a in c(1:l)){
    l2 <- length(AA[[a]])
    for(b in c(1:l2)){
      value <- AA[[a]][b]
      if(value > initialcodon && value < endcodon){
        AAs[a,(value%/%4 - (del-1))] <- "F"
      }
    }
  }
AA <- gregexpr("(\\sTT[AG])|(\\sCT[GACT])",codons,useBytes = FALSE)
  l <- length(AA)
```

```
  for(a in c(1:l)){
    l2 <- length(AA[[a]])
    for(b in c(1:l2)){
      value <- AA[[a]][b]
      if(value > initialcodon && value < endcodon){
        AAs[a,(value%/%4 - (del-1))] <- "L"
      }
    }
  }
AA <- gregexpr("(\\sTC[GCAT])|(\\sAG[TC])",codons,useBytes = FALSE)
  l <- length(AA)
  for(a in c(1:l)){
    l2 <- length(AA[[a]])
    for(b in c(1:l2)){
      value <- AA[[a]][b]
      if(value > initialcodon && value < endcodon){
        AAs[a,(value%/%4 - (del-1))] <- "S"
      }
    }
  }
AA <- gregexpr("\\sTA[TC]",codons,useBytes = FALSE)
  l <- length(AA)
  for(a in c(1:l)){
    l2 <- length(AA[[a]])
    for(b in c(1:l2)){
      value <- AA[[a]][b]
      if(value > initialcodon && value < endcodon){
        AAs[a,(value%/%4 - (del-1))] <- "Y"
      }
    }
  }
AA <- gregexpr("\\sTAG",codons,useBytes = FALSE)
  l <- length(AA)
  for(a in c(1:l)){
    l2 <- length(AA[[a]])
    for(b in c(1:l2)){
      value <- AA[[a]][b]
      if(value > initialcodon && value < endcodon){
        AAs[a,(value%/%4 - (del-1))] <- "TAG"
      }
    }
  }
AA <- gregexpr("\\sTAA",codons,useBytes = FALSE)
  l <- length(AA)
```

```
  for(a in c(1:l)){
    l2 <- length(AA[[a]])
    for(b in c(1:l2)){
      value <- AA[[a]][b]
      if(value > initialcodon && value < endcodon){
        AAs[a,(value%/%4 - (del-1))] <- NA
      }
    }
  }
AA <- gregexpr("\\sTG[TC]",codons,useBytes = FALSE)
  l <- length(AA)
  for(a in c(1:l)){
    l2 <- length(AA[[a]])
    for(b in c(1:l2)){
      value <- AA[[a]][b]
      if(value > initialcodon && value < endcodon){
        AAs[a,(value%/%4 - (del-1))] <- "C"
      }
    }
  }
AA <- gregexpr("\\sTGA",codons,useBytes = FALSE)
  l <- length(AA)
  for(a in c(1:l)){
    l2 <- length(AA[[a]])
    for(b in c(1:l2)){
      value <- AA[[a]][b]
      if(value > initialcodon && value < endcodon){
        AAs[a,(value%/%4 - (del-1))] <- NA
      }
    }
  }
AA <- gregexpr("\\sTGG",codons,useBytes = FALSE)
  l <- length(AA)
  for(a in c(1:l)){
    l2 <- length(AA[[a]])
    for(b in c(1:l2)){
      value <- AA[[a]][b]
      if(value > initialcodon && value < endcodon){
        AAs[a,(value%/%4 - (del-1))] <- "W"
      }
    }
  }
AA <- gregexpr("\\sCC[GCAT]",codons,useBytes = FALSE)
  l <- length(AA)
```

```
  for(a in c(1:l)){
    l2 <- length(AA[[a]])
    for(b in c(1:l2)){
      value <- AA[[a]][b]
      if(value > initialcodon && value < endcodon){
        AAs[a,(value%/%4 - (del-1))] <- "P"
      }
    }
  }
AA <- gregexpr("\\sCA[CT]",codons,useBytes = FALSE)
  l <- length(AA)
  for(a in c(1:l)){
    l2 <- length(AA[[a]])
    for(b in c(1:l2)){
      value <- AA[[a]][b]
      if(value > initialcodon && value < endcodon){
        AAs[a,(value%/%4 - (del-1))] <- "H"
      }
    }
  }
AA <- gregexpr("\\sCA[AG]",codons,useBytes = FALSE)
  l <- length(AA)
  for(a in c(1:l)){
    l2 <- length(AA[[a]])
    for(b in c(1:l2)){
      value <- AA[[a]][b]
      if(value > initialcodon && value < endcodon){
        AAs[a,(value%/%4 - (del-1))] <- "Q"
      }
    }
  }
AA <- gregexpr("(\\sCG[GCAT])|(\\sAG[GA])",codons,useBytes = FALSE)
  l <- length(AA)
  for(a in c(1:l)){
    l2 <- length(AA[[a]])
    for(b in c(1:l2)){
      value <- AA[[a]][b]
      if(value > initialcodon && value < endcodon){
        AAs[a,(value%/%4 - (del-1))] <- "R"
      }
    }
  }
AA <- gregexpr("\\sAT[CAT]",codons,useBytes = FALSE)
  l <- length(AA)
```

```
    for(a in c(1:l)){
      l2 <- length(AA[[a]])
      for(b in c(1:l2)){
        value <- AA[[a]][b]
        if(value > initialcodon && value < endcodon){
          AAs[a,(value%/%4 - (del-1))] <- "I"
        }
      }
    }
  AA <- gregexpr("\\sATG",codons,useBytes = FALSE)
    l <- length(AA)
    for(a in c(1:l)){
      l2 <- length(AA[[a]])
      for(b in c(1:l2)){
        value <- AA[[a]][b]
        if(value > initialcodon && value < endcodon){
          AAs[a,(value%/%4 - (del-1))] <- "M"
        }
      }
    }
  AA <- gregexpr("\\sAC[GCAT]",codons,useBytes = FALSE)
    l <- length(AA)
    for(a in c(1:l)){
      l2 <- length(AA[[a]])
      for(b in c(1:l2)){
        value <- AA[[a]][b]
        if(value > initialcodon && value < endcodon){
          AAs[a,(value%/%4 - (del-1))] <- "T"
        }
      }
    }
  AA <- gregexpr("\\sAA[CT]",codons,useBytes = FALSE)
    l <- length(AA)
    for(a in c(1:l)){
      l2 <- length(AA[[a]])
      for(b in c(1:l2)){
        value <- AA[[a]][b]
        if(value > initialcodon && value < endcodon){
          AAs[a,(value%/%4 - (del-1))] <- "N"
        }
      }
    }
  AA <- gregexpr("\\sAA[AG]",codons,useBytes = FALSE)
    l <- length(AA)
```

```
  for(a in c(1:l)){
    l2 <- length(AA[[a]])
    for(b in c(1:l2)){
      value <- AA[[a]][b]
      if(value > initialcodon && value < endcodon){
        AAs[a,(value%/%4 - (del-1))] <- "K"
      }
    }
  }
AA <- gregexpr("\\sGT[GACT]",codons,useBytes = FALSE)
  l <- length(AA)
  for(a in c(1:l)){
    l2 <- length(AA[[a]])
    for(b in c(1:l2)){
      value <- AA[[a]][b]
      if(value > initialcodon && value < endcodon){
        AAs[a,(value%/%4 - (del-1))] <- "V"
      }
    }
  }
AA <- gregexpr("\\sGC[GACT]",codons,useBytes = FALSE)
  l <- length(AA)
  for(a in c(1:l)){
    l2 <- length(AA[[a]])
    for(b in c(1:l2)){
      value <- AA[[a]][b]
      if(value > initialcodon && value < endcodon){
        AAs[a,(value%/%4 - (del-1))] <- "A"
      }
    }
  }
AA <- gregexpr("\\sGA[TC]",codons,useBytes = FALSE)
  l <- length(AA)
  for(a in c(1:l)){
    l2 <- length(AA[[a]])
    for(b in c(1:l2)){
      value <- AA[[a]][b]
      if(value > initialcodon && value < endcodon){
        AAs[a,(value%/%4 - (del-1))] <- "D"
      }
    }
  }
AA <- gregexpr("\\sGA[AG]",codons,useBytes = FALSE)
  l <- length(AA)
```

```
  for(a in c(1:l)){
    l2 <- length(AA[[a]])
    for(b in c(1:l2)){
      value <- AA[[a]][b]
      if(value > initialcodon && value < endcodon){
        AAs[a,(value%/%4 - (del-1))] <- "E"
      }
    }
  }
AA <- gregexpr("\\sGG[GACT]",codons,useBytes = FALSE)
  l <- length(AA)
  for(a in c(1:l)){
    l2 <- length(AA[[a]])
    for(b in c(1:l2)){
      value <- AA[[a]][b]
      if(value > initialcodon && value < endcodon){
        AAs[a,(value%/%4 - (del-1))] <- "G"
      }
    }
  }
AAs <- as.data.frame(AAs)
#this gives unique amino acid sequences
UniqueAAs <- AAs %>% group_by_all() %>% count()
UniqueAAs <- UniqueAAs[order(-UniqueAAs$n),]
UniqueAAs <- UniqueAAs[apply(UniqueAAs,1,function(row) all(row !=
 0)),]
UniqueAAs <- na.omit(UniqueAAs)
UniqueAAsR2pp <- UniqueAAs

#this counts sequences that have TAG codons, sequences that have more
 than one are only counted once
TAGreg <- regexpr("\\sTAG",codons)
TAGtable <- table(TAGreg)
percentTAG <-
 sum(TAGtable[2:length(TAGtable)])/length(codons)*100#percent of
 sequences containing TAG

#this creates a matrix of amino acid sequences that do not contain TAG
 codons
TAGpos <- which(AAs == "TAG")
TAGrow <- TAGpos%%nrow(AAs)
AAsnoTAG <- AAs[-TAGrow,]
```

```
#this creates heatmap for amino acid frequency per library position,
 change scale according to values
AAtable <- apply(AAs,2,function(x) table(factor(x,levels=aa)))
AAtable <- as.matrix(AAtable/length(codons))
colnames(AAtable) <- c(1:lib)
heatmapcolors <- colorRampPalette(brewer.pal(9,"Blues"))(100)
sc <- seq(0.0,0.6,by=0.006)
AAheatmap <- heatmap.2(AAtable, Rowv = NA, Colv = NA, col =
 heatmapcolors, density.info = "none", scale = "none", trace = "none",
 breaks = sc,  xlab = "Position in Library", ylab = "Codon", margins =
 c(3,4), dendrogram = "none")

#this creates projected heatmap based on NNK randomized codons
randomAAs <- matrix(0,21,lib,dimnames =
 list(rownames(AAtable),c(1:lib)))
randomAAs[c("A","G","P","T","V"),] <- 2/32
randomAAs[c("C","H","Q","N","K","Y","D","E","W","I","M","TAG","F"),]
 <- 1/32
randomAAs[c("L","S","R"),] <- 3/32
NNKheatmap <- heatmap.2(randomAAs, Rowv = NA, Colv = NA, col =
 heatmapcolors, density.info = "none", scale = "none", trace = "none",
 breaks = sc,  xlab = "Position in Library", ylab = "Codon", margins =
 c(3,4), dendrogram = "none")

#this creates heatmap showing bias from random, change scale with
 respect to range of values
lscale <- seq(-1,4,by=5/100)
librarybias <- (AAtable - randomAAs)/randomAAs
Biasheatmap <- heatmap.2(librarybias, Rowv = NA, Colv = NA, col =
 heatmapcolors, density.info = "none", scale = "none", trace = "none",
 breaks = lscale,  xlab = "Position in Library", ylab = "Codon",
 margins = c(3,4), dendrogram = "none")
librarybias <- as.data.frame(librarybias)

#this creates heatmap for AAsnoTAG
AAnoTAGtable <- apply(AAsnoTAG,2,function(x)
 table(factor(x,levels=aa)))
AAnoTAGtable <- as.matrix(AAnoTAGtable/nrow(AAsnoTAG))
colnames(AAnoTAGtable) <- c(1:lib)
heatmapcolors <- colorRampPalette(brewer.pal(9,"Blues"))(100)
sc <- seq(0.0,0.3,by=0.003)
AAheatmap <- heatmap.2(AAnoTAGtable, Rowv = NA, Colv = NA, col =
 heatmapcolors, density.info = "none", scale = "none", trace = "none",
```

```
breaks = sc,  xlab = "Position in Library", ylab = "Codon", margins =
c(3,4), dendrogram = "none")

#writes csv files for uniqueAAs and bias heatmaps change path to make
file
path <- "/Users/traehampton/Documents/Research/Sequencing Results/Next
Gen Sequencing/21510Wns_N21170"
write.csv(UniqueAAs, paste(path,"/12mernegnegUniqueAAs.csv", sep =
""), row.names = F)
write.csv(TAGtable, paste(path,"/12mernegnegTAGtable.csv", sep = ""),
row.names = T)
write.csv(librarybias, paste(path,"/12mernegnegLibraryBias.csv", sep =
""), row.names = T)
```

**Supplementary Script 2: Script for Volcano plots and Enrichment data**

```
#this gives overall enrichment of each peptide sequence by comparing
Round 1 and Round 4 Library
library(prodlim)
path <- "/Users/traehampton/Documents/Research/Sequencing Results/Next
Gen Sequencing/22468Wns_N22179"
lib <- 14
match <-
row.match(as.data.frame(UniqueAAsAlloc[,1:lib]),as.data.frame(UniqueAA
sBoc[,1:lib]))
matchseq <- which(is.na(match)==FALSE)
percentenriched <-
(UniqueAAsBoc[match[matchseq],lib+1]/sum(UniqueAAsBoc[,lib+1])-
UniqueAAsAlloc[matchseq,lib+1]/sum(UniqueAAsAlloc$n))/(UniqueAAsAlloc[
matchseq,lib+1]/sum(UniqueAAsAlloc$n))
enrichedseq <- UniqueAAsBoc[match[matchseq],]
enrichedseq$enrichment <- percentenriched[,1]
enrichedseq <- enrichedseq[order(-enrichedseq$enrichment),]
enrichedseq <- as.data.frame(enrichedseq)
UniqueAAsAlloc$percent <- UniqueAAsAlloc$n/sum(UniqueAAsAlloc$n)*100
UniqueAAsBoc$percent <- UniqueAAsBoc$n/sum(UniqueAAsBoc$n)*100
#Change names to whatever you want
write.csv(enrichedseq, paste(path,"/28R1vR3.csv", sep = ""), row.names
= F)
write.csv(UniqueAAsAlloc, paste(path,"/AllocCombined.csv", sep = ""),
row.names = F)
write.csv(UniqueAAsBoc, paste(path,"/BocCombined.csv", sep = ""),
row.names = F)
```

```
#this is for volcano plot if the sequencing is done in more than one
 replicate
path <- "/Users/traehampton/Documents/Research/Sequencing Results/Next
 Gen Sequencing/22081Wns_N22028"
lib <- 14
UniqueAAsA1R3$percent <-
 UniqueAAsA1R3[,lib+1]/sum(UniqueAAsA1R3[,lib+1])*100
UniqueAAsA2R3$percent <-
 UniqueAAsA2R3[,lib+1]/sum(UniqueAAsA2R3[,lib+1])*100
match1 <-
 row.match(as.data.frame(UniqueAAsA1R3[,1:lib]),as.data.frame(UniqueAAs
 A2R3[,1:lib]))
match2 <-
 row.match(as.data.frame(UniqueAAsA2R3[,1:lib]),as.data.frame(UniqueAAs
 A1R3[,1:lib]))
na1 <-which(is.na(match1) == TRUE)
na2 <-which(is.na(match2) == TRUE)
UniqueAAsA1R3$percentA2 <- 0
UniqueAAsA2R3$percentA2 <- 0
UniqueAAsA1R3[,lib+3] <- UniqueAAsA2R3[match1,lib+2]
UniqueAAsA1R3 <- as.data.frame(UniqueAAsA1R3)
UniqueAAsA1R3[which(is.na(UniqueAAsA1R3$percentA2)==TRUE),lib+3] <- 0
UniqueAAsAlloc <- rbind(UniqueAAsA1R3,UniqueAAsA2R3[na2,])
UniqueAAsAlloc$average <- apply(UniqueAAsAlloc[,(lib+2):(lib+3)], 1,
 mean)
UniqueAAsAlloc$sd <- apply(UniqueAAsAlloc[,(lib+2):(lib+3)], 1, sd)

UniqueAAsA1R1$percent <-
 UniqueAAsA1R1[,lib+1]/sum(UniqueAAsA1R1[,lib+1])*100
UniqueAAsA2R1$percent <-
 UniqueAAsA2R1[,lib+1]/sum(UniqueAAsA2R1[,lib+1])*100
match1 <-
 row.match(as.data.frame(UniqueAAsA1R1[,1:lib]),as.data.frame(UniqueAAs
 A2R1[,1:lib]))
match2 <-
 row.match(as.data.frame(UniqueAAsA2R1[,1:lib]),as.data.frame(UniqueAAs
 A1R1[,1:lib]))
na1 <-which(is.na(match1) == TRUE)
na2 <-which(is.na(match2) == TRUE)
UniqueAAsA1R1$percentA2 <- 0
UniqueAAsA2R1$percentA2 <- 0
UniqueAAsA1R1[,lib+3] <- UniqueAAsA2R1[match1,lib+2]
UniqueAAsA1R1 <- as.data.frame(UniqueAAsA1R1)
UniqueAAsA1R1[which(is.na(UniqueAAsA1R1$percentA2)==TRUE),lib+3] <- 0
```

```
UniqueAAsAllocR1 <- rbind(UniqueAAsA1R1,UniqueAAsA2R1[na2,])
UniqueAAsAllocR1$average <- apply(UniqueAAsAllocR1[,(lib+2):(lib+3)],
 1, mean)
UniqueAAsAllocR1$sd <- apply(UniqueAAsAllocR1[,(lib+2):(lib+3)], 1,
 sd)

match1 <-
 row.match(as.data.frame(UniqueAAsAlloc[,1:lib]),as.data.frame(UniqueAA
 sAllocR1[,1:lib]))
match2 <-
 row.match(as.data.frame(UniqueAAsAllocR1[,1:lib]),as.data.frame(Unique
 AAsAlloc[,1:lib]))
AllocR1vR3 <- UniqueAAsAlloc
AllocR1vR3$AllocR1percent1 <- UniqueAAsAllocR1[match1,lib+2]
AllocR1vR3$AllocR1percent2 <- UniqueAAsAllocR1[match1,lib+3]
AllocR1vR3$Allocaverage <- UniqueAAsAllocR1[match1,lib+4]
AllocR1vR3 <- AllocR1vR3[-
 which(is.na(AllocR1vR3$AllocR1percent2)==TRUE),]

AllocR1vR3$neglog10pvalue <- 0
AllocR1vR3$log2foldchange <- 0
for(a in c(1:nrow(AllocR1vR3))){
  t <- t.test(AllocR1vR3[a,(lib+2):(lib+3)],
 AllocR1vR3[a,(lib+6):(lib+7)])
  AllocR1vR3[a,"neglog10pvalue"] <- -log(t[[3]])
  if(sum(AllocR1vR3[a,(lib+6):(lib+7)]) != 0){
    AllocR1vR3[a,"log2foldchange"] <- log2(t[[5]][1]/t[[5]][2])
  }
  else{
    AllocR1vR3[a,"log2foldchange"] <- 20
  }
}

plot(AllocR1vR3$log2foldchange, AllocR1vR3$neglog10pvalue)

rows <- c(1:lib,lib+9,lib+10)
write.csv(AllocR1vR3[,rows], paste(path,"/AllocR1vR3Volcanoplot.csv",
 sep = ""), row.names = T)

#copy for boc enrichment volcano plot
path <- "/Users/traehampton/Documents/Research/Sequencing Results/Next
 Gen Sequencing/22081Wns_N22028"
lib <- 14
```

```
UniqueAAsB1R3$percent <-
 UniqueAAsB1R3[,lib+1]/sum(UniqueAAsB1R3[,lib+1])*100
UniqueAAsB2R3$percent <-
 UniqueAAsB2R3[,lib+1]/sum(UniqueAAsB2R3[,lib+1])*100
match1 <-
 row.match(as.data.frame(UniqueAAsB1R3[,1:lib]),as.data.frame(UniqueAAs
 B2R3[,1:lib]))
match2 <-
 row.match(as.data.frame(UniqueAAsB2R3[,1:lib]),as.data.frame(UniqueAAs
 B1R3[,1:lib]))
na1 <-which(is.na(match1) == TRUE)
na2 <-which(is.na(match2) == TRUE)
UniqueAAsB1R3$percentA2 <- 0
UniqueAAsB2R3$percentA2 <- 0
UniqueAAsB1R3[,lib+3] <- UniqueAAsB2R3[match1,lib+2]
UniqueAAsB1R3 <- as.data.frame(UniqueAAsB1R3)
UniqueAAsB1R3[which(is.na(UniqueAAsB1R3$percentA2)==TRUE),lib+3] <- 0
UniqueAAsBoc <- rbind(UniqueAAsB1R3,UniqueAAsB2R3[na2,])
UniqueAAsBoc$average <- apply(UniqueAAsBoc[,(lib+2):(lib+3)], 1, mean)
UniqueAAsBoc$sd <- apply(UniqueAAsBoc[,(lib+2):(lib+3)], 1, sd)

UniqueAAsB1R1$percent <-
 UniqueAAsB1R1[,lib+1]/sum(UniqueAAsB1R1[,lib+1])*100
UniqueAAsB2R1$percent <-
 UniqueAAsB2R1[,lib+1]/sum(UniqueAAsB2R1[,lib+1])*100
match1 <-
 row.match(as.data.frame(UniqueAAsB1R1[,1:lib]),as.data.frame(UniqueAAs
 B2R1[,1:lib]))
match2 <-
 row.match(as.data.frame(UniqueAAsB2R1[,1:lib]),as.data.frame(UniqueAAs
 B1R1[,1:lib]))
na1 <-which(is.na(match1) == TRUE)
na2 <-which(is.na(match2) == TRUE)
UniqueAAsB1R1$percentA2 <- 0
UniqueAAsB2R1$percentA2 <- 0
UniqueAAsB1R1[,lib+3] <- UniqueAAsB2R1[match1,lib+2]
UniqueAAsB1R1 <- as.data.frame(UniqueAAsB1R1)
UniqueAAsB1R1[which(is.na(UniqueAAsB1R1$percentA2)==TRUE),lib+3] <- 0
UniqueAAsBocR1 <- rbind(UniqueAAsB1R1,UniqueAAsB2R1[na2,])
UniqueAAsBocR1$average <- apply(UniqueAAsBocR1[,(lib+2):(lib+3)], 1,
 mean)
UniqueAAsBocR1$sd <- apply(UniqueAAsBocR1[,(lib+2):(lib+3)], 1, sd)
```

```
match1 <-
 row.match(as.data.frame(UniqueAAsBoc[,1:lib]),as.data.frame(UniqueAAsB
 ocR1[,1:lib]))
match2 <-
 row.match(as.data.frame(UniqueAAsBocR1[,1:lib]),as.data.frame(UniqueAA
 sBoc[,1:lib]))
BocR1vR3 <- UniqueAAsBoc
BocR1vR3$BocR1percent1 <- UniqueAAsBocR1[match1,lib+2]
BocR1vR3$BocR1percent2 <- UniqueAAsBocR1[match1,lib+3]
BocR1vR3$BocR1average <- UniqueAAsBocR1[match1,lib+4]
BocR1vR3 <- BocR1vR3[-which(is.na(BocR1vR3$BocR1percent2)==TRUE),]

BocR1vR3$neglog10pvalue <- 0
BocR1vR3$log2foldchange <- 0
for(a in c(1:nrow(BocR1vR3))){
  t <- t.test(BocR1vR3[a,(lib+2):(lib+3)],
 BocR1vR3[a,(lib+6):(lib+7)])
  BocR1vR3[a,"neglog10pvalue"] <- -log(t[[3]])
  if(sum(BocR1vR3[a,(lib+6):(lib+7)]) != 0){
    BocR1vR3[a,"log2foldchange"] <- log2(t[[5]][1]/t[[5]][2])
  }
  else{
    BocR1vR3[a,"log2foldchange"] <- 20
  }
}

plot(BocR1vR3$log2foldchange, BocR1vR3$neglog10pvalue)

rows <- c(1:lib,lib+9,lib+10)
write.csv(BocR1vR3[,rows], paste(path,"/BocR1vR3Volcanoplot.csv", sep
 = ""), row.names = T)

#Compares top100 R3 sequences and percentages shared between Boc and
 Alloc
match1 <-
 row.match(as.data.frame(UniqueAAsBoc[1:100,1:lib]),as.data.frame(Uniqu
 eAAsAlloc[,1:lib]))
PercentBocinAlloc <- sum(na.omit(UniqueAAsAlloc[match1,lib+4]))
match2 <-
 row.match(as.data.frame(UniqueAAsAlloc[1:100,1:lib]),as.data.frame(Uni
 queAAsBoc[,1:lib]))
PercentAllocinBoc <- sum(na.omit(UniqueAAsBoc[match2,lib+4]))
```
